# Innate immune pathways act synergistically to constrain RNA virus evolution in *Drosophila melanogaster*

**DOI:** 10.1101/2021.07.16.452470

**Authors:** Vanesa Mongelli, Sebastian Lequime, Athanasios Kousathanas, Valérie Gausson, Hervé Blanc, Lluis Quintana-Murci, Santiago F. Elena, Maria-Carla Saleh

## Abstract

Host-pathogen interactions impose recurrent selective pressures that lead to constant adaptation and counter-adaptation in both competing species. Here, we sought to study this evolutionary arms-race and assessed the impact of the innate immune system on viral population diversity and evolution, using *D. melanogaster* as model host and its natural pathogen Drosophila C virus (DCV). We first isogenized eight fly genotypes generating animals defective for RNAi, Imd and Toll innate immune pathways and also pathogen sensing and gut renewal pathways. Wild-type or mutant flies were then orally infected and DCV was serially passaged ten times. Viral population diversity was studied after each viral passage by high-throughput sequencing, and infection phenotypes were assessed at the beginning and at the end of the passaging scheme. We found that the absence of any of the various immune pathways studied increased viral genetic diversity and attenuated the viruses. Strikingly, these effects were observed in both host factors with antiviral properties and host factors with antibacterial properties. Together, our results indicate that the innate immunity system as a whole, and not specific antiviral defense pathways in isolation, generally constrains viral diversity and evolution.

## Introduction

Interaction between hosts and pathogens trigger defense and counter-defense mechanisms that often result in reciprocal adaptation and coevolution of both organisms^1^. Empirical evidence of such arms-race involving both species can be drawn from genome-wide analysis from hosts and pathogens and in experimental evolution settings. For example, evolutionary analysis of mammalian genomes have revealed evidence of host-virus coevolution between different retroviruses and antiviral factors^2,3^, and in plants, host resistance genes and virulence genes encoded by pathogens have been found to co-evolve^4^. Likewise, between bacteria and their infecting bacteriophages, experimental co-evolution studies resulted in the occurrence of genetic variants in both a bacterial lipopolysaccharide synthesis gene and the phage tail fiber gene which binds to lipopolysaccharide during adsorption^5^. In nematodes and their pathogenic bacteria, the number of toxin-expressing plasmids varies during adaptation to the host^6^.

In insects, analysis of sequences within and between drosophila species showed evidence of adaptive evolution in immunity related genes^7–10^. In a study using mosquitoes, West Nile virus, and siRNAs deep sequencing, it was found that the regions of the viral genome more intensively targeted by the RNA interference (RNAi) mechanism contained a higher number of mutations than viral genome regions less affected by this pathway, suggesting that this antiviral defense mechanism imposes a selective pressure to the viral population^11^. Similar observations on the selective pressure imposed by the RNAi pathway on viral evolution have been done in plants and human infecting viruses^12–16^. *Drosophila melanogaster* is a well-studied insect model to decipher virus-host interactions and therefore the impact of the host antiviral immunity on viral diversity and evolution. Different drosophila immunity pathways and mechanisms are involved in antiviral defense^17,18^. As is the case for all invertebrates, defense against pathogens in drosophila relies on innate immunity, which constitutes not only the first, but the exclusive defense against microbes. Innate immunity is characterized by the recognition of pathogen derived molecules, called pathogen-associated molecular patterns (PAMPs), by host encoded receptors (pathogen recognition receptors - PRRs), which leads to a rapid defense response.

The RNAi mechanism is known to play a central role in drosophila antiviral defense, mainly through the action of the small interfering (si) RNA pathway^19–22^. Antiviral RNAi is triggered by virtually any insect-infecting virus, targeting its genome in a sequence specific manner to control infection. Several other pathways have antiviral properties in flies, but their roles in defense against virus seem to be virus specific. The Toll and Imd (Immune deficiency) pathways, originally described to be involved in antibacterial and antifungal responses, have been shown to play a role in antiviral defense against Drosophila C virus (DCV), Cricket paralysis virus (CrPV), Drosophila X virus, Nora virus, and Flock house virus^23–26^. The Janus kinase signal transducers and activators of transcription (Jak-STAT) pathway can be activated upon DCV and CrPV infections in flies, triggering the expression of antiviral factors^27,28^.

DCV, a positive sense single stranded RNA virus from the genus *Cripavirus* within the *Dicistrioviridae* family and *Picornavirales* order^29^, is a well characterized natural pathogen of the fruit fly that can be found in laboratory and wild populations^30^. As for many other drosophila-infecting viruses, defense against DCV depends on the joint action of different innate immune pathways and mechanisms. RNAi, Toll and Imd pathways, but also the gene Vago, play a role in the defense against this virus^20,24–27,31–33^. DCV is thought to be acquired by ingestion in natural conditions^30,34,35^. For orally acquired pathogens, the digestive tract, and the gut in particular, represents the first host defense barrier. Despite many studies using oral bacterial infections^36^, the role of gut-specific antiviral responses in drosophila is not fully understood. Gut triggered responses against bacterial pathogens include the production of reactive oxygen species (ROS), antimicrobial peptides, and also tissue repair and regeneration mechanisms. Furthermore, the maintenance of gut homeostasis after tissue damage caused by pathogenic bacteria relies on the activity of JAK-STAT and epidermal growth factor receptor (EGFR) pathways, amongst others^37–39^. In the hallmark of viral infections, a role of the Imd and ERK pathways in the antiviral response in the gut has been suggested^24,40^. It is important to note that, like many other RNA viruses with error-prone polymerases and fast replication kinetics, DCV exists as large populations composed of a cloud of genetically related mutant variants, a phenomenon known as viral quasispecies or mutant swarm^41^. Viral quasispecies constitute a dynamic repertoire of genetic and phenotypic variability that renders great adaptability.

In this work, we leveraged the vast knowledge on antiviral mechanisms and extensive genetic tool-box available for *D. melanogaster*, the intrinsic variability of DCV mutant swarm, and the high depth power of next generation sequencing, to study the impact of innate immunity pathways on viral diversity and evolution. We aimed to determine not only if each pathway has a specific impact on the selective pressure imposed to DCV mutant swarms, but also their relative impact. In addition, we investigated possible links between selected viral variants (viral function) and specific defense mechanisms. Our results show that the host genotype has an impact on viral genetic diversity regardless of the immune pathway being affected and this is accompanied by an increase in survival of infected flies along evolutionary passages. We also describe complex mutation dynamics, with several examples of clonal interference in which increases in frequency of adaptive mutations have been displaced by other mutations of stronger effect that arose in different genetic backgrounds. Overall, our results highlight that innate immunity pathways constrain RNA virus evolution and further demonstrate that antiviral responses in drosophila are likely polygenic.

## Results

### Production of fly mutant lines for innate immune pathways

To reduce genetic variation due to differences in genetic background, mutant flies defective on the RNAi, Imd and Toll immunity pathways and also pathogen sensing and gut renewal mechanism, were isogenized prior to beginning viral evolution studies. Homozygous loss-of-function lines for Argonaute-2 (*Ago-2^414^*), Dicer-2 (*Dcr-2^L811fsX^* and *Dcr-2^R416X^*), Dorsal-related immunity factor (*Dif^1^*), Relish (*Rel^E20^*), Spätzle (*spz^2^*), and Vago (*Vago^ΔM10^*) and hippomorphic mutant line for Epidermal growth factor receptor (*Egfr^t1^*) were produced in the same genetic background by crossing parental lines at least 10 times to *w^1118^* flies. Infection phenotypes of the newly produced fly lines were characterized by following their survival after inoculation of DCV by intrathoracic injection (Supplementary Figure 1a). As previously described, *Dcr-2^L811fsX/L811fsX^, Dcr-2^R416X/R416X^* and *Ago-2^414/414^* mutants infected with DCV succumbed faster than *w^1118^* flies^20,21^, as well as *Vago^ΔM10/ΔM10^*mutants^33^. Toll pathway mutants *spz^2/2^* and *Dif^1/1^* and Imd pathway mutant *Rel^E20/E20^* were less sensitive to DCV infection than *w^1118^* flies as they died later than *w^1118^* flies (Supplementary Figure 1a); however, these mutants kept the increased susceptibility to infection by Gram + and Gram − bacteria respectively (Supplementary Figure 1b and 1c). No difference in virus-induced mortality was found between *w^1118^* and *Egfr^t1/t1^* mutant flies (Supplementary Figure 1a). This set of isogenic mutant flies with contrasting phenotypes to DCV infection provided us with the host model system to perform the experimental viral evolution assay.

### Experimental DCV evolution

To study the impact of innate immune pathways on virus population diversity and evolution, DCV was serially passaged in *w^1118^* flies and in the isogenic innate immunity deficient fly lines (Figure 1a). DCV population diversity was studied after each passage by next generation sequencing (NGS) and DCV virulence was analyzed at the beginning and at the end of the evolution experiment.

**Figure 1.**
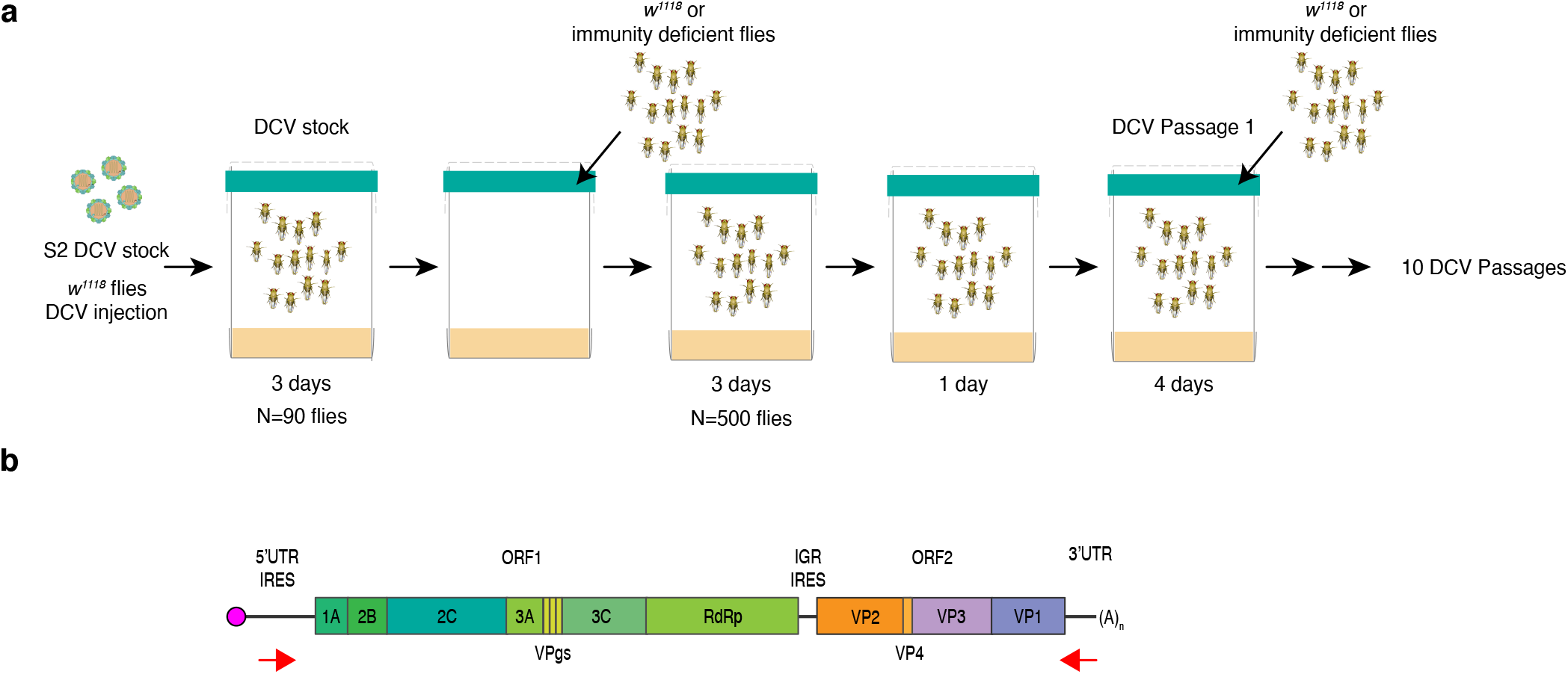
Experimental design. **a)** Scheme of the experimental DCV evolution assay. To produce the starting DCV stock (DCV stock) 5 to 6 days old *w^1118^* were intrathoracically injected with 100 TCID_50_ units of DCV from a stock produced in S2 drosophila cells (S2 DCV stock) or mock infected. At 4 dpi, N = 90 DCV infected flies were placed in cages containing fresh drosophila medium, left during 3 days and then removed to place in these DCV contaminated or mock contaminated cages N = 500 5 to 6 days old *w^1118^* or mutant flies (males and females). Flies were allowed to feed *ad libitum* during 3 days (oral inoculation period), then moved to a clean cage for 1 day, and further placed into a new clean cage and left during 4 days, when they were harvested (DCV passage 1, P = 1). Contaminated cages were used to place a new group of 500 flies. This procedure was repeated 10 times (10 DCV passages, P = 1 to P = 10) and replicated twice (biological replicates BR1 and BR2). For each passage, and genotype, half of the harvested flies were used to PCR-amplify the complete DCV genome followed by deep-sequenced. The other half was used to produce viral stock for passages P = 1 and P = 10 from each genotype for phenotypic characterization. **b)** Scheme of DCV genome and the localization of primers used to amplify the complete viral genome. The viral genome is composed of single-stranded positive-sense RNA and encodes for two ORFs which are transcribed as polyproteins. The first ORF (ORF 1) encodes for the non-structural viral proteins, 1A: viral silencing suppressor, 2C: RNA helicase, VPg: viral genome-linked protein, 3C: protease, RdRp: RNA-dependent RNA polymerase, 2B and 3A: are thought to be involved in the assembly of the viral replication complex. The second ORF (ORF 2) encodes for DCV structural proteins VP1 to VP4 which constitute the viral capsid.

To follow viral infection during the course of the experiment, viral load was determined by TCID_50_ and prevalence (percentage of flies positive for TCID_50_) was calculated for all passages in individual flies from DCV contaminated cages. We found that for most fly genotypes and for both biological replicates, 60% or more of the flies became infected with DCV along the 10 viral passages (Supplementary Figure 2a). When studying viral loads across passages only *w^1118^, Ago-2^414/414^* and *Rel^E20/E20^* fly lines displayed large variability while viral load in the other fly genotypes remained relatively stable along passages (Supplementary Figure 2b).

To assess the impact that fly genotype, biological replicate, and viral passage has on viral loads, the log-transformed TCID_50_ values from Supplementary Figure 2b were fitted to the generalized linear model (GLM) described in the Materials and Methods section. In short, the model incorporates fly genotype and experimental block as orthogonal factors and passage as covariable. Highly significant differences were observed on viral load among fly genotypes (test of the intercept: *χ*^2^ = 146.734, 8 d.f., *p* < 0.001) that were of very large magnitude 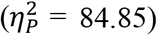, thus confirming that DCV load strongly varies among host genotypes. A significant effect was also observed for the viral passages (test of the covariable: *χ*^2^ = 5.075, 1 d.f., *p* = 0.024), indicating overall differences in viral accumulation among passages, though the magnitude of this effect was rather small 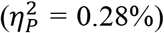. Regarding second-order interactions among factors and the covariable, a significant interaction exist between fly genotype and experimental block (*χ*^2^ = 27.082, 8 d.f., *p* < 0.001) indicating that some of the differences observed in virus accumulation among host genotypes differed among biological replicates, and between fly genotype and evolutionary passage (*χ*^2^ = 52.511, 8 d.f., *p* < 0.001). However, despite being statistically significant, these two effects were of very small magnitude (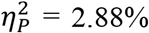 and 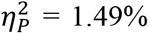, respectively) and likely biologically irrelevant. Likewise, the third-order interaction was statistically significant (*χ*^2^ = 86.023, 8 d.f., *p* < 0.001), suggesting that the differences in viral load among experimental blocks observed for a particular host genotype also depended on the evolutionary passages, although once again the effect could be considered as minor 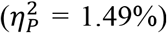. Next, we evaluated whether differences exist in viral load between immune competent (*w^1118^*) and the different mutant fly genotypes. In all eight cases, DCV accumulated to significantly higher levels in the immune deficient flies than in the wild-type ones (*p* < 0.001), with the smallest significant difference corresponding to viral populations replicating in *Rel^E20/E20^* and *Dif^1/1^* and the largest to those replicating in *Egfr^t1/t1^* and *Dcr-2^R416X/R416X^* (Supplementary Figure 2c).

Overall, these results show that in both immune competent (*w^1118^*) and immune deficient flies, DCV oral infection was maintained along passages and confirm that mutant flies are more permissive to DCV infection.

### Viral nucleotide diversity differently evolves in each host genotype

To look into the selective pressure imposed by the drosophila innate immune pathways on DCV population dynamics, we analyzed virus genome diversity after each passage. Half of the population of infected flies was used to sequence DCV full-length genome by NGS (Figure 1a and 1b). The viral stocks used to start the experiment, S2 DCV stock and DCV stock, were also sequenced. Sequencing analysis was performed using the computational pipeline Viral Variance Analysis (ViVan)^42^. Sequence coverage was at least 8,000 reads per position on the genome. To determine the error rate of the sequencing procedure, including library preparation, four sequencing technical replicates from S2 DCV stock were used (Supplementary Figure 3). A frequency threshold of 0.0028 was used for all subsequent analyses based on variant detection and frequency correlation between technical replicates (see Methods section).

To determine if the lack of activity of a given innate immunity pathway had an impact on viral population genetic diversity, we calculated the site-averaged nucleotide diversity (π) on all polymorphic sites (*n* = 1869) across the full-length viral genome, and in the different DCV genomic regions, present in the full dataset (all passages, including the S2 DCV stock). We compared the viral nucleotide diversity present in each fly genotype to each other (Table 1 and Supplementary Table 1) and fly genotypes were sorted in four groups according to their increasing viral nucleotide diversity: group 1 (less diversity): *w^1118^, Dcr-2^L811fs/[L811fsX^* and *Dif^1/1^* fly lines; group 2: *Dif^1/1^, Dcr-2^L811fs/[L811fsX^, Rel^E20/E20^, spz^2/2^*, and *Dcr-2^R416X/R416X^* fly lines; group 3: *Dcr-2^L811fs/[L811fsX^, Rel^E20/E20^, spz^2/2^, Dcr-2^R416X/R416X^*, and *Ago-2^414/414^* fly lines; group 4 (more diversity): containing *spz^2/2^, Dcr-2^R416X/R416X^, Ago-2^414/414^, Egfr^t1/t1^*, and *Vago^ΔM10/ΔM10^* fly lines (Figure 2, Table 1 and Supplementary Table 1). Next, we analyzed the trajectories of viral nucleotide diversity along passages and determined if the host genotype, viral passages, biological replicate, and the interactions between these factors had an impact on the evolution of diversity (Figure 2 and Table 2). We observed that only the fly genotype had a statistically significant impact on the differences in π found for each host genotype (*χ*^2^ = 25.545, 8 d.f., *p* = 0.001) (Table 2).

**Figure 2.**
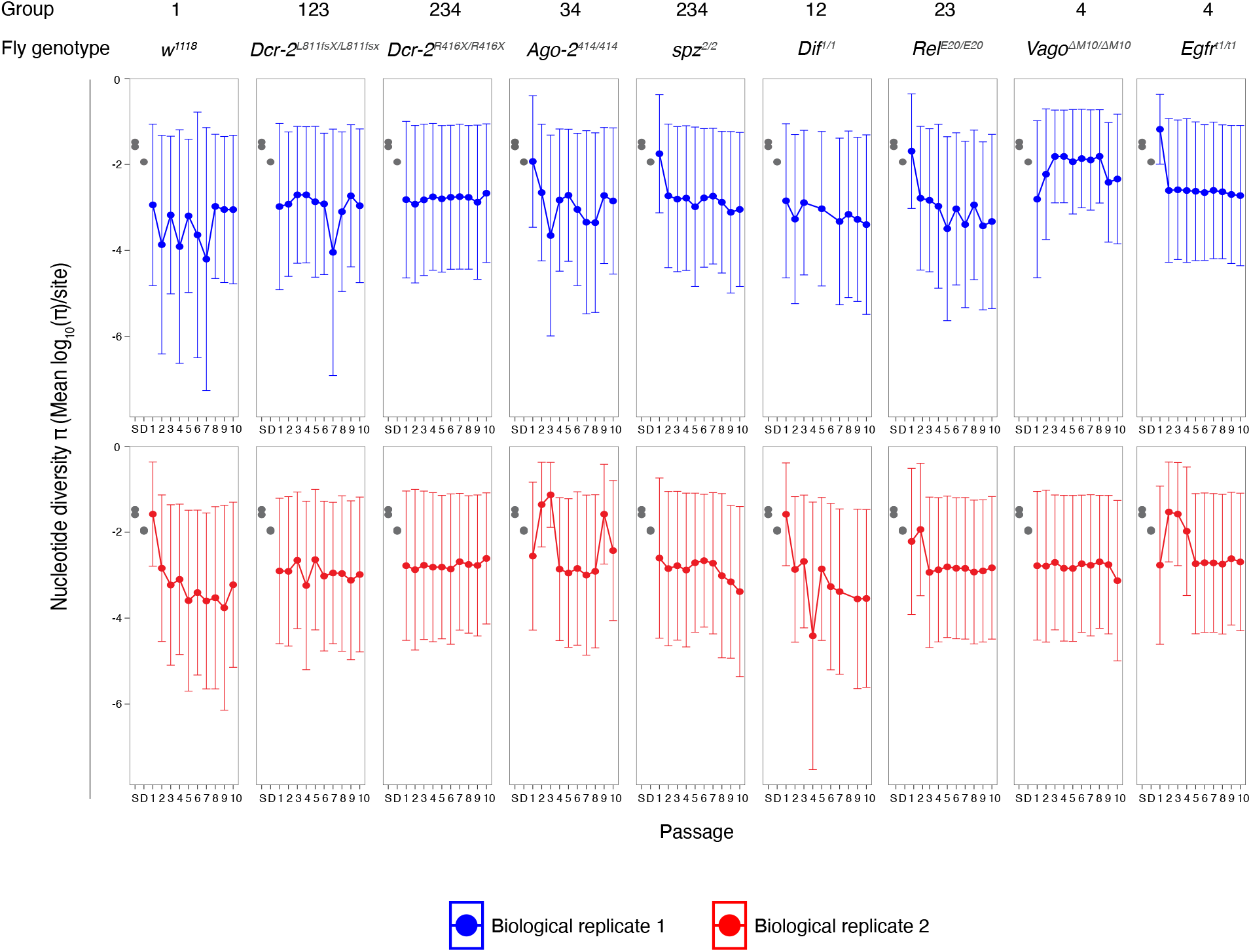
Viral nucleotide diversity differently evolves in each host genotype. Trajectory of the site-averaged nucleotide diversity (π) on all polymorphic sites (n = 1869) across the full-length viral genome, and in the different DCV genomic regions.

**Table 1.**
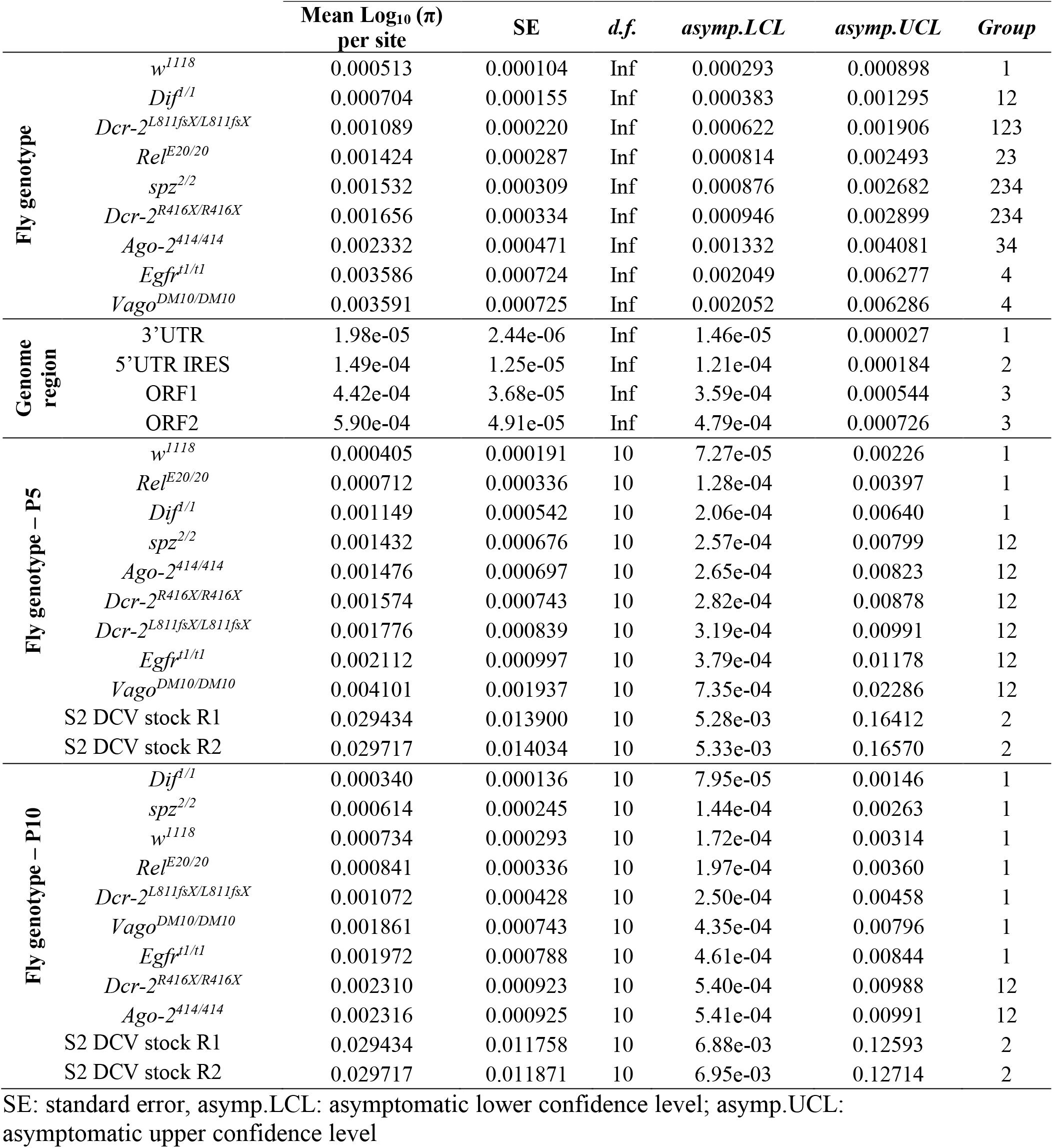
Comparison of viral nucleotide diversities (π) by Bonferroni *post hoc* test based on pairwise comparisons from Supplementary Table 1.

**Table 2.** Analysis of the impact of each experimental variable on the evolution of viral nucleotide diversity (π) considering the full-length viral genome and all viral passages.

| Source of variation | LR $\chi^2$ | *d.f.* | Pr(>$\chi^2$) |
| --- | --- | --- | --- |
| BR | 2.2528 | 1 | 0.133372 |
| P | 1.6460 | 1 | 0.199498 |
| G | 25.5447 | 8 | 0.001256 ** |
| (BR)xP | 0.0024 | 1 | 0.960572 |
| (BR)xG | 14.2963 | 8 | 0.074361 |
| PxG | 12.1679 | 8 | 0.143867 |
| (BR)xPxG | 10.4253 | 8 | 0.236435 |

BR: Biological replicate; P: viral passage; G: fly genotype
LR: likelihood ratio $\chi^2$ ; Pr(>Chisq): p-value

We then wondered if the general differences observed in viral nucleotide diversity, between fly genotypes, were associated to a particular viral genomic region (*i.e*., if a determined viral function was affected during the evolution experiment). To do so, we used the π values on all polymorphic sites (*n* = 1869) across the full-length viral genome and present in the full database. Of note, IGR IRES was not included because its lack of genetic variation prevented us from determining its nucleotide diversity value. Pairwise comparisons of the viral genetic diversity within the genomic regions allowed us to distinguish three main groups: group 1 (less diversity): 3’UTR; group 2: 5’UTR IRES; and group 3 (more diversity): ORF1 and ORF2 (Table 1 and Supplementary Table 1). We found that the fly genotype had a statistically significant effect in π (*χ*^2^ = 27.178, 8 d.f., *p* < 0.001), as well as specific viral genomic regions (*χ*^2^ = 11.698, 8 d.f., *p* = 0.008). As a second-order interaction an effect of the fly genotype and the biological replicate was found (*χ*^2^ = 16.314, 8 d.f., *p* = 0.038).

To determine how the virus evolved from the starting viral stock (S2 DCV stock) in each fly genotype, viral nucleotide diversity in *P* = 1, *P* = 5 and *P* =10 was subsequently compared with the diversity in the S2 DCV stock. Pairwise comparison between viral nucleotide diversity in each fly genotypes in *P* = 1 versus S2 DCV stock, yield no statistically significant difference (p = 1.000) (Supplementary Table 1). In *P* = 5 viral diversity was reduced only in *w^1118^* (p = 0.0042 and p = 0.0041), *Rel^E20/E20^* (p = 0.0130 and p = 0.0128), and *Dif^1/1^* (p = 0.0366 and p = 0.0358) mutants when compared to the starting viral stock (Group 1 – Table 1 and Supplementary Table 1). In *P* = 10 nucleotide diversity present in all fly genotypes (p < 0.05), except for *Dcr-2^R416X/R416X^* (p = 0.0624 and p = 0.0608) and *Ago-2^414/414^* (p = 0.0628 and p = 0.0612) mutant lines, was reduced compared to S2 DCV stock (Group 1 – Table 1 and Supplementary Table 1). We then performed an ANOVA analysis to test whether biological replicates and fly genotype could explain the difference in nucleotide diversity of the evolved strain versus the S2 DCV stock. We found no significant impact on diversity in *P* = 1 (biological replicate: *χ*^2^ = 0.0479, 10 d.f., *p* = 0.1313 and fly genotype: *χ*^2^ = 4.5369, 10 d.f., *p* = 0.3682. However, in both *P* = 5 and 10, the fly genotype but not the biological replicate was found to have a significant impact (*χ*^2^ = 7.119, 10 d.f., *p* = 0.002 and *χ*^2^ = 8.010, 10 d.f., *p* < 0.001 respectively).

These results indicate that viral nucleotide diversity differently evolved in each host genotype, with coding regions of the virus displaying higher levels of nucleotide diversity than non-coding regions. They also indicate a general decrease in viral population diversity, independently of the fly genotype, when compared to the starting viral stock.

### Viral population diversity derives from preexisting standing genetic variation

Next, we examined if the levels of viral diversity observed in DCV populations from innate immunity mutants compared to the *w^1118^* were accompanied with the fixation of particular genetic changes in the mutant swarms, and whether (*i*) these changes can be associated to fitness effects and (*ii*) potentially adaptive mutations arose in response to particular immune responses. To do so, we estimated the selection coefficients for each SNPs using their variation in frequency across evolutionary time (Figure 3 and Supplementary Figure 4), using a classic population genetics approach^43^ (Table 3). Thirty-six SNPs yielded significant estimates of selection coefficients (this number reduces to 10 if a stricter FDR correction is applied; Table 3). Twenty-one of them were already detected in the ancestral S2 DCV stock, henceforth a maximum of 15 new SNPs might have arisen during the evolution experiment. Estimated selection coefficients for all these SNPs ranged between −0.304 per passage (synonymous mutation RdRp/C5713U) and 1.204 per passage (VP2/G6311C nonsynonymous change R16P), with a median value of 0.286 per passage (interquartile rank = 0.265). Nine mutations were observed in more than one lineage (range 2 - 7 times), with synonymous mutations VP3/U7824C appearing in seven lineages of six different host genotypes and mutation 5’UTR/A280U in five lineages of five host genotypes (Table 3). These nine SNPs were all present in the S2 DCV stock. Indeed, the frequency of SNPs among evolving lineages is significantly correlated with their frequency in the ancestral S2 DCV stock (Pearson’s *r* = 0.401, 36 df, *p* = 0.013), but not with their measured fitness effect (*r* = −0.091, 36 df, *p* = 0.588).

**Figure 3.**
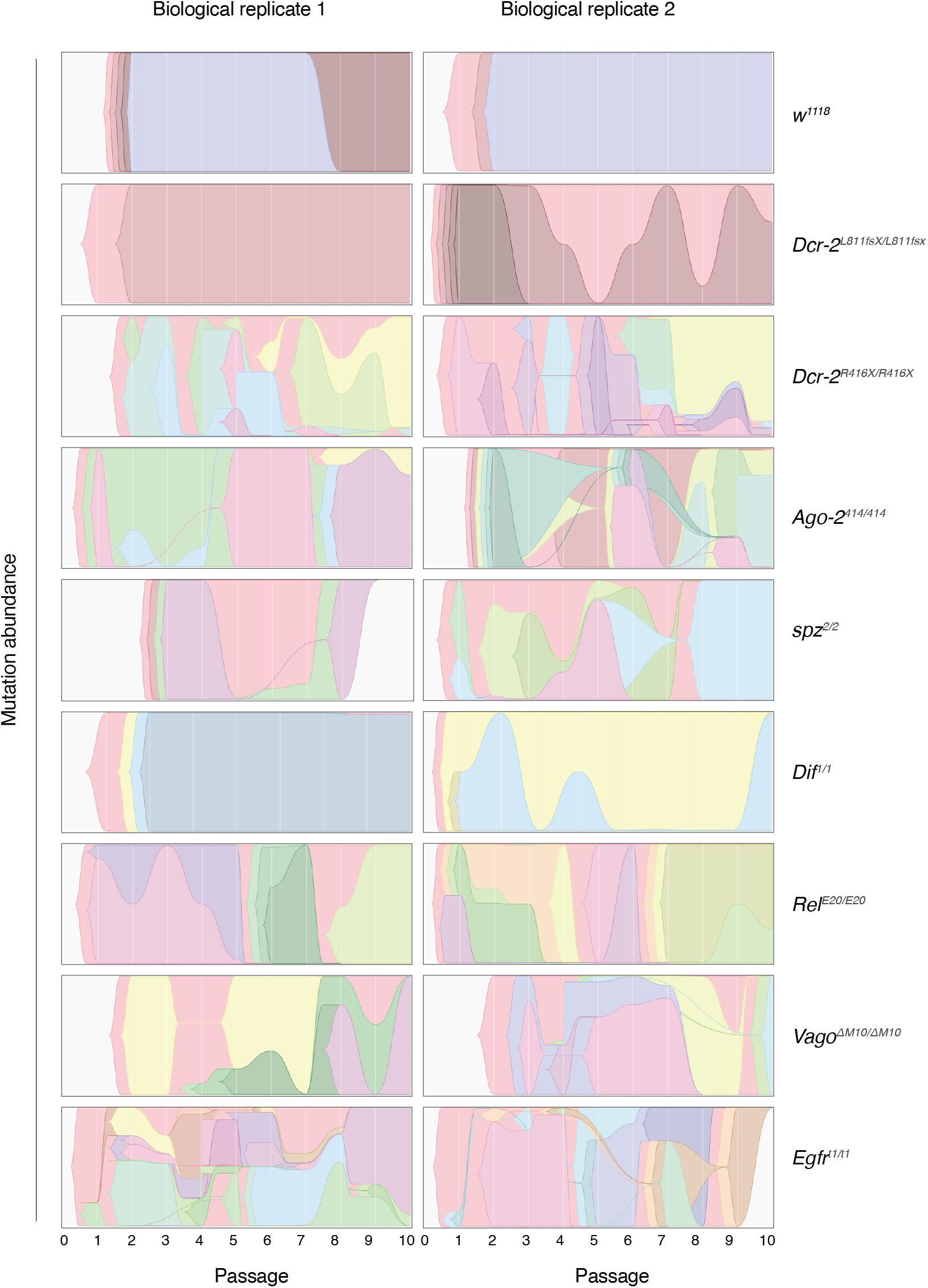
Trajectories of DCV variants across passages. Muller plots illustrating the dynamics of SNPs’ frequencies along evolutionary time.

**Table 3.**
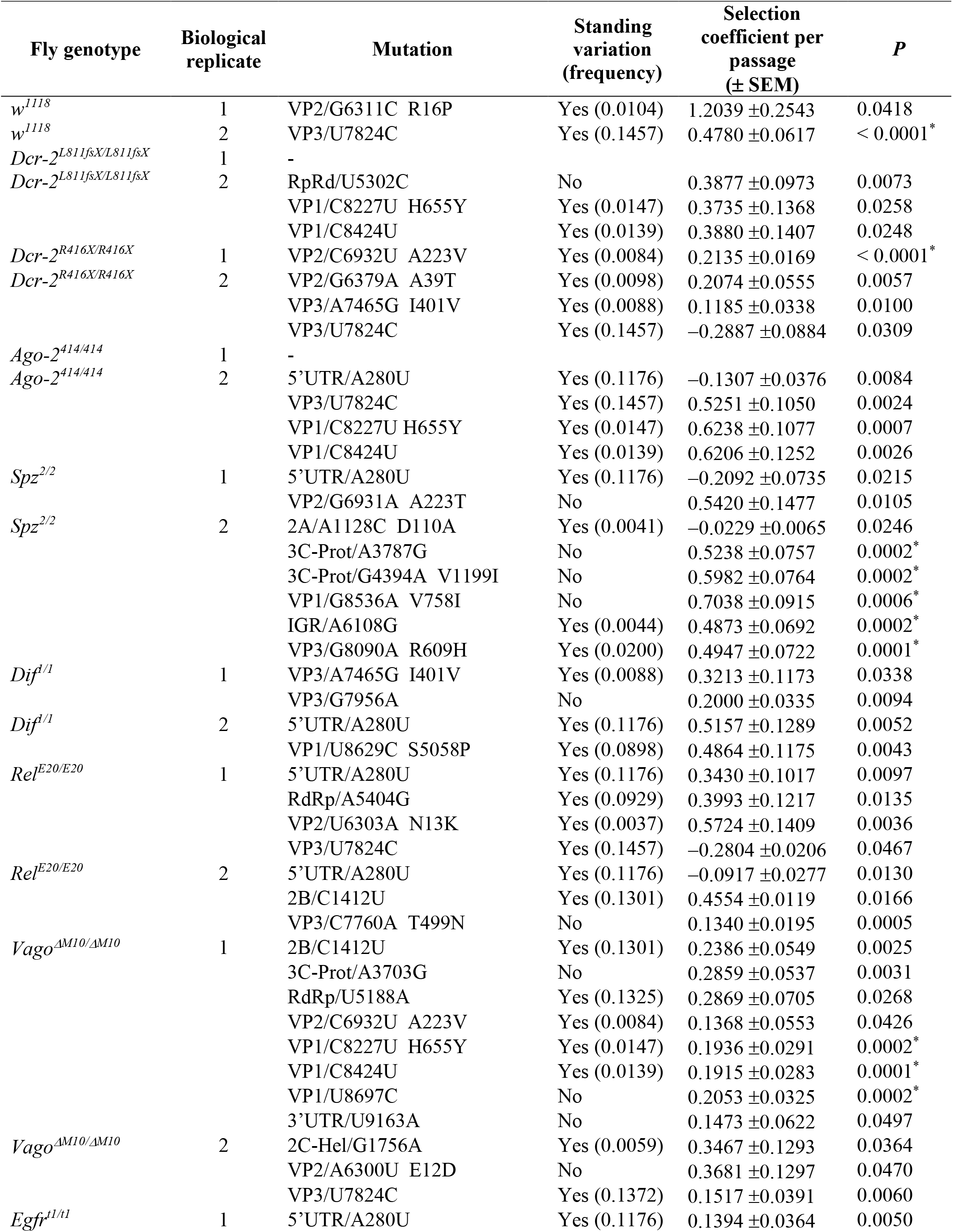
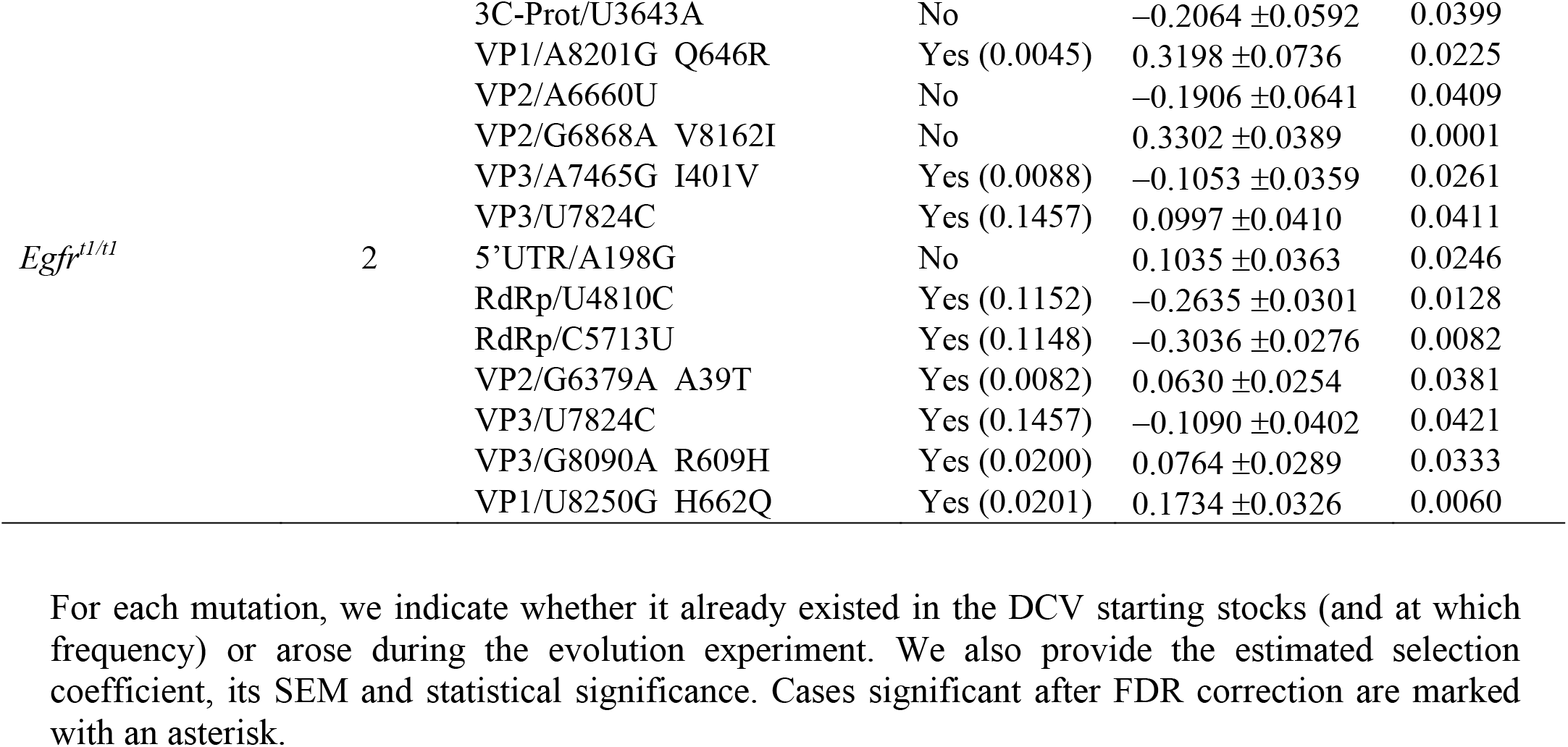
Mutations for which significant estimates of fitness effects have been obtained.

An interesting question is whether the fitness effects associated to each of these nine SNPs were the same across all genotypes or, conversely, fitness effects were host genotype-dependent. To test this hypothesis, we performed one-way ANOVA tests comparing fitness effects (Table 3) across the corresponding host genotypes. In all cases, significant differences were observed (*F* ≥ 15.637 and *p* ≤ 0.001, and ≥ 93.99% of total observed variance in fitness effects explained by true genetic differences among host genotypes), supporting the notion that fitness effects are indeed host-genotype dependent. A pertinent example is the case of the synonymous mutation VP3/U7824C, which was the most prevalent mutation (*F*_6,45_ = 158.862, *p* < 0.001, 99.37% of genetic variance). In this case, a *post hoc* Bonferroni test shows that host genotypes can be classified into three groups according to the fitness effect of this SNP. In genotypes *Dcr-2^R416X/R416X^* and *Rel^E20/E20^*, the mutation has a deleterious effect (on average, −0.2260 per passage); in genotypes *Egfr^t1/t1^* and *Vago^ΔM10/ΔM10^*, the mutation is moderately beneficial (on average, 0.1257 per passage; and in genotypes *w^1118^* and *Ago-2^414/414^*, the mutation had a strong beneficial effect (on average, 0.502 per passage).

As shown in Figure 3 and Supplementary Figure 4a, some SNPs show a strong parallelism in their temporal dynamics, suggesting they might be linked into haplotypes. This is particularly relevant for mutations shown in Table 5. To test this possibility, we computed all pairwise Pearson correlation coefficients between mutations’ frequencies along evolutionary time. The results of these analyses are shown in Supplementary Figure 4b to 4k as heatmaps. Again, as an illustrative example, we discuss here the case of the viral population BR2 evolved in *Ago-2^414/414^* (Supplementary Figure 4d). Synonymous mutations VP3/U7824C and VP1/C8424U and nonsynonymous mutation VP1/C8227U (H655Y) are all linked into the same haplotype (*r* ≥ 0.998, *p* < 0.001). Since these three mutations already existed in the S2 DCV stock, it is conceivable that the haplotype already existed and has been selected as a unit. Indeed, the fitness effects estimated for these three mutations are undistinguishable (one-way ANOVA: *F*_2, 22_ = 1.781, *p* = 0.192; average fitness effect 0.590 ±0.032 per passage) thus suggesting that the estimated value corresponds to the haplotype as a unit. The absence of this haplotype in *Ago-2^414/414^* BR1 suggests it was lost during the transmission bottleneck from S2 cells to flies. Interestingly, mutations VP1/C8424U VP1/C8227U appear also linked into the same haplotype in population BR2 evolved in *Dcr-2^L811fsX/L811fsX^* (Supplementary Figure 4b). These two cases, as well as populations BR1 evolved in *Rel^E20/E20^*, BR2 evolved in *spz^2/2^* and BR1 and BR2 evolved in *Vago^ΔM10/ΔM10^* illustrate easy to interpret haplotypes (Supplementary Figure 4f, 4e, 4h, and 4i). Other viral populations, especially those evolved in *Egfr^t1/t1^* flies, show much more complex patterns (Supplementary Figure 4j and 4k) in which haplotypes change along time by acquiring *de novo* mutations.

When mapping the 36 SNPs found to have significant estimates of selection coefficients in the viral genome (Figure 4), we found that two mapped to the 5’UTR IRES, twelve to the ORF1 coding for the non-structural proteins of the virus, one to the viral IGR IRES, 20 mutations were in DCV ORF2 encoding for the viral structural proteins and one mutation mapped to the 3’UTR. From the nine mutations observed in ORF1, four mapped to the 3C viral protease and five to the RdRp, and only one of those mutations in the 3C protein was non-synonymous. From the 20 mutations from ORF2, eight mapped to VP2, five to VP3 and seven to VP1, the three major DCV predicted capsid proteins.

**Figure 4.**
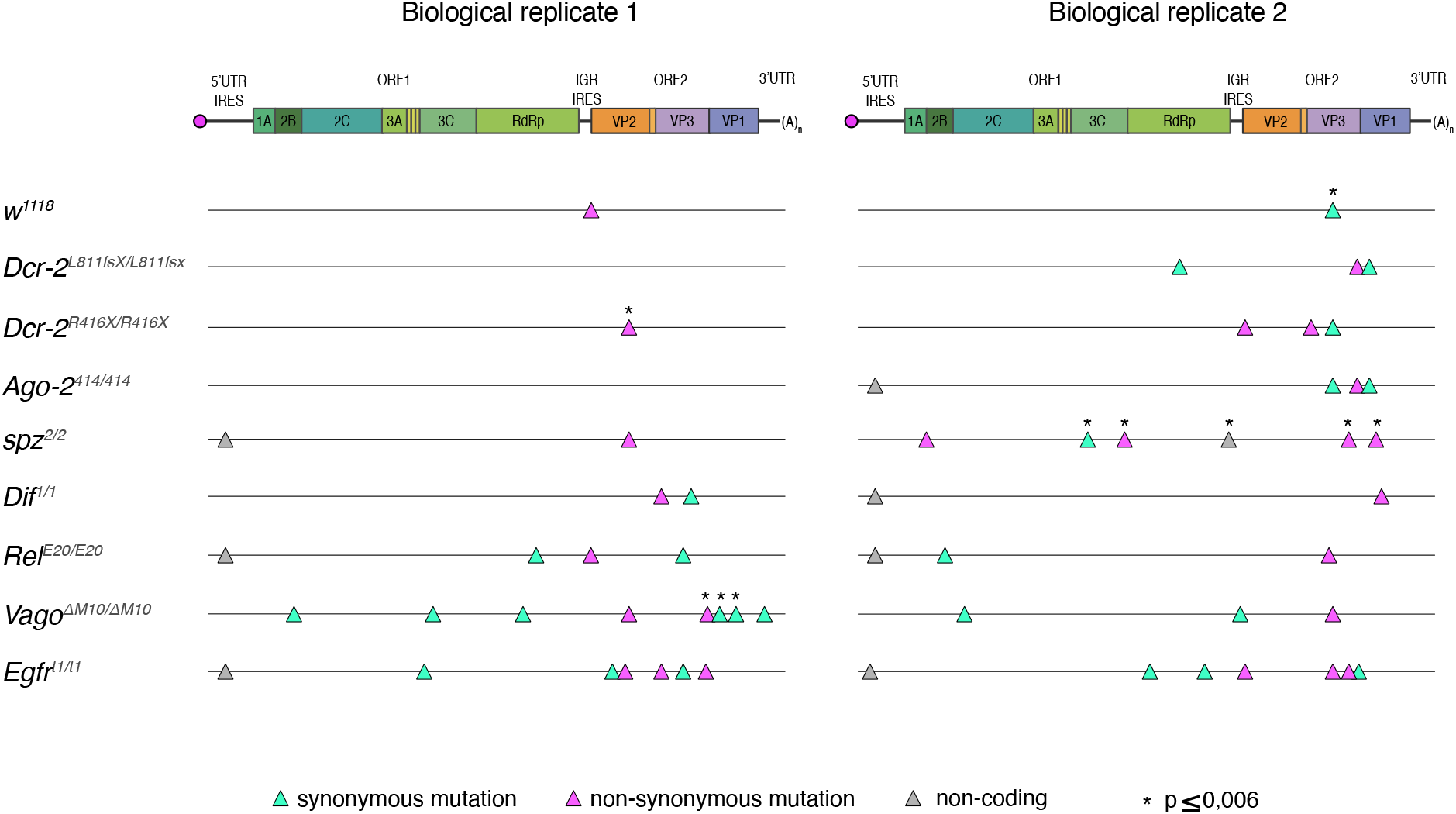
SNPs for which significant estimates of fitness effects have been obtained mapped on the viral genome. Green triangles represent synonymous mutations, pink triangles nonsynonymous mutations and gray triangles mutations in non-coding sequences. Cases significant after FDR correction are marked with an asterisk.

Taken together, these results show that viral population diversity mainly derived from preexisting standing genetic variation in the ancestral DCV population. Furthermore, temporal dynamics of population diversity is linked to the fly genotype in which the virus evolved.

### DCV virulence is not affected by the absence of immune pathways

Finally, we wondered if DCV virulence varied among each lineage in the different fly genotypes. Infectious DCV stocks were produced from viral passages *P* = 1 and *P* = 10 and from all fly genotypes. Because DCV is not lethal during oral infection^31^, we intrathoracically inoculated *w^1118^* flies with 10 TCID_50_ of DCV stocks derived from *P* = 1 or *P* = 10 and from each fly genotype. Survival of the flies was determined daily. We found that *w^1118^* flies were less sensitive to viral infection when inoculated with DCV stocks derived from *P* = 10 since they succumbed later than those inoculated with stocks from *P* = 1 for most DCV stock origins (Figure 5a and Supplementary Table 2). Notable exceptions were DCV stocks from BR2 of *Vago^ΔM10/ΔM10^* mutant flies, for which *w^1118^* flies were more sensitive to *P* = 10 than to *P* = 1, and stocks from BR1 of *spz^2/2^* and BR2 of *Egfr^t1/t1^* mutant flies, for which no difference in survival after infection with DCV between *P* = 1 and *P* = 10 was detected.

**Figure 5.**
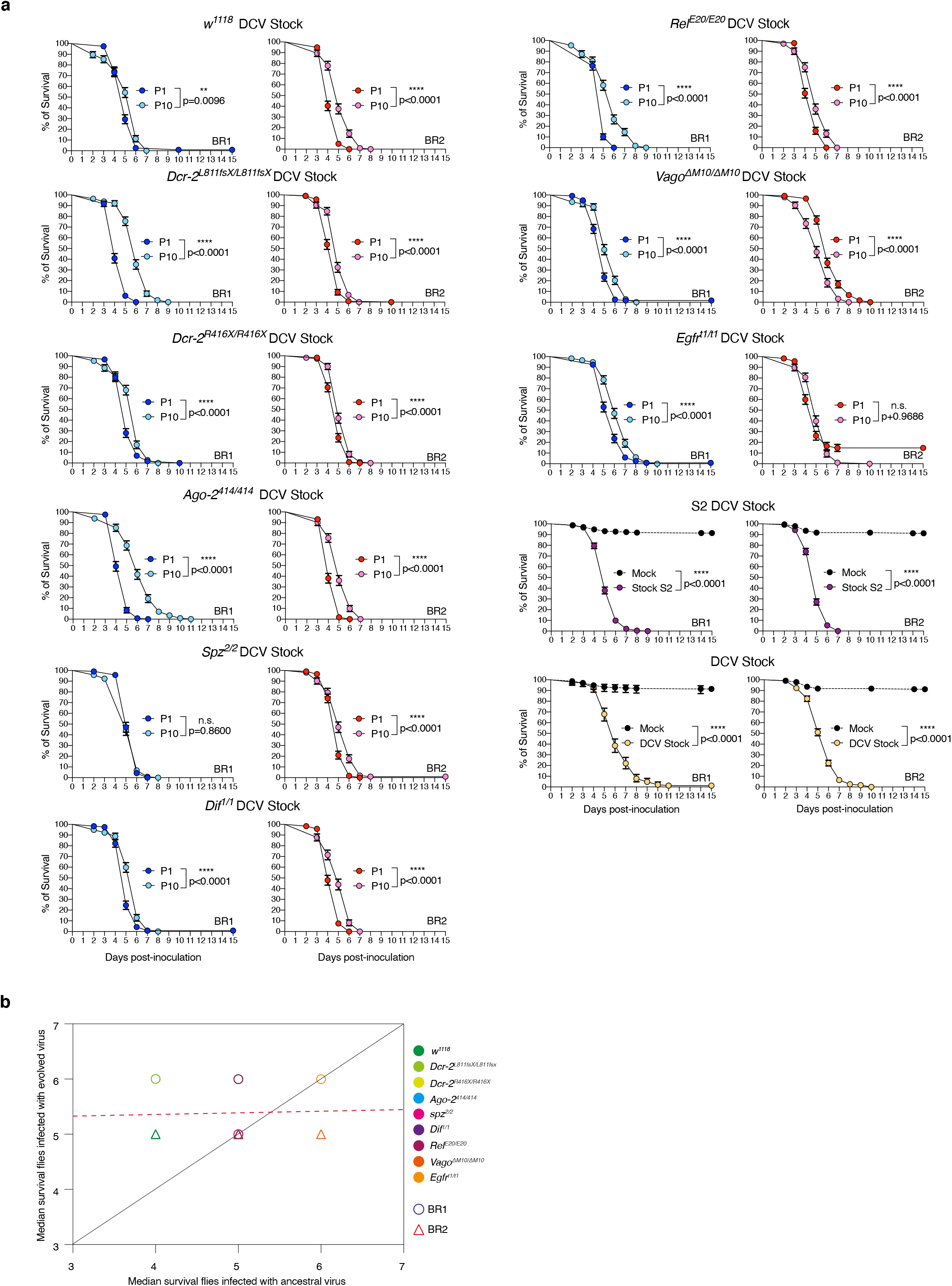
DCV virulence is not affected by the absence of immune pathways. DCV infectious stocks were prepared from viral passages P = 1 and P = 10 and from each fly genotype. *w^1118^* flies were intrathoracically inoculated with 10 TCID_50_ units of each DCV stock and survival of the flies was measured daily. **a)** Survival curves shown in the figure are the combination of the two independent replicates, with three technical replicates each, of a total of at least N = 98 flies per treatment. Error bars indicate +/-1 SEM; n.s., not significant. Survival curves were compared via log-rank (Mantel–Cox) tests. **b)** Test of the contribution of historical contingency evolved (P = 10) vs ancestral (P = 1) DCV virulence. The black line represents the linear regression and the dashed red line represents the expected relationship under the null hypothesis of ancestral differences in DCV virulence which are maintained after evolution despite noise introduced by random events (mutation and drift).

A fundamental question in evolutionary biology is the role that past evolutionary events may have in the outcome of evolution^44^. If ongoing evolution is strongly contingent with past evolutionary events, ancestral phenotypic differences should be retained to some extent, while if other evolutionary forces such as selection and stochastic events (*i.e*., mutation and genetic drift) dominate, then ancestral differences can be eroded and, in the extreme case, even fully removed. Here, we observed significant differences in the performance of the ancestral DCV across the eight host genotypes. To test whether these differences are still observable in the evolved population, we compared the median survival time (Figure 5a and Supplementary Table 2) for DCV populations isolated at the beginning of the evolution experiment *P* = 1 and at the end *P* = 10 (Figure 5b). Under the null hypothesis of strong historical contingency, it is expected that data will fit to a regression line of slope 1 and intercepting the ordinate axis at 0. However, if ancestral differences have been removed, data would fit significantly better to a regression line with a slope smaller than one and with an intercept greater than zero^44^. Figure 5b shows the data and its fit to the null hypothesis (solid black line) and the alternative hypothesis (dashed red line). A partial *F*-test shows that adding an intercept to the regression equation significantly improves the fit (*F*_1,16_ = 28.437, *p* < 0.001), thus supporting the notion that ancestral differences among host genotypes have been removed by the action of subsequent adaptation, that is, the fixation of beneficial mutations.

## Discussion

In this work we aimed at determining the overall impact of innate immunity on viral evolution. Based on the arms-race hypothesis, we speculated that if a given host defense mechanism imposes a specific selective pressure on a particular pathogen function, the absence of this defense mechanism would result in the relaxation of the selective constraint, which would be in turn detectable in the pathogen at the genomic and phenotypic levels. We found that viral population diversity evolved differently according to each fly genotype; however, a reduction of ancestral genetic variation, regardless of the immune pathway affected, was also observed. Our results indicate that antiviral responses are polygenic; there is not a specific, main immune defense mechanism against a particular virus, but instead a repertoire of defense mechanisms that are triggered after infection and that might interact with each other.

Our results are compatible with a pervasive presence of clonal interference. In the absence of sexual reproduction, clonal interference is the process by which beneficial alleles originated in different clades within a population compete to each other, resulting in one of them reaching fixation. Subsequently, the outcompeted beneficial allele may appear in the new dominant genetic background and, assuming no negative epistasis among both loci, become fixed. As a consequence, beneficial mutations may fix sequentially, thus slowing down the rate of adaptation^45^. Given their large effective population size and high mutation rates, viral populations are expected to contain considerable amounts of potentially beneficial standing variation, making them prone to clonal interference. Indeed, it has been previously shown to operate in experimental populations of vesicular stomatitis virus adapting to cell cultures^46,47^, in bacteriophage φX174 populations adapting to harsh saline environments^48^, in tobacco etch virus adapting to novel plant host species^49^, among HIV-1 escape variants within individual patients^50^, and also at the epidemiological level among influenza A virus lineages diversifying antigenically^51^. In our own results, clonal interference can be observed in populations BR1 evolved in *Dcr-2^L811fsX/L811fsX^*, BR1 evolved in *Ago-2^414/414^*, BR1 evolved in *spz^2/2^*, BR2 evolved in *Rel^E20/E20^*, and BR2 evolved in *Vago^ΔM10/ΔM10^* all share similar patterns in which some beneficial allele (or haplotypes) rose in frequency, reached a peak at some intermediate passage, then declined in frequency and were finally outcompeted by a different beneficial mutation (or haplotype) that had lower initial frequency. For example, the nonsynonymous mutation VP2/G6931A (A223T) appeared *de novo* in population BR1 evolved in *spz*^2/2^, and outcompeted several mutations likely linked in a haplotype (Figure 3). Tightly linked to clonal interference is the concept of leap-frogging^52^, in which the beneficial mutation that ends up dominating the population is less genetically related to the previously dominant haplotype than to the common ancestor of both (Figure 3). The VP2/G6931A mutation well illustrates this example, as it appeared in a genetic background that was minoritarian rather than in the dominant one. Likewise, the mutation VP2/G6311C (R16P), observed in BR1 evolved in *w^1118^* flies, appeared in a low frequency genetic background different from the most abundant one in previous passages. Finally, the haplotype containing five different mutations observed in BR2 evolved in *spz^2/2^* became dominant in frequency after *P* = 6, outcompeting two other mutations that were dominating the population until then.

The existence and fixation of haplotypes along our evolution experiment deserves further discussion. Linked mutations generate three possible interference effects^53^. Firstly, all mutations might contribute additively, or may be involved in positive epistasis, to the fitness of the haplotype as a whole, thus increasing its chances to become fixed. Secondly, hitchhiking and genetic draft may occur, by which deleterious or neutral alleles are driven to fixation along with a linked beneficial allele. Thirdly, there may be background selection by which the spread of a beneficial allele is impeded, or at least delayed, owing to the presence of linked deleterious alleles. For instance, we can hypothesize that haplotype VP3/U7824C-VP1/C8227U-VP1/C8424U, which swept to fixation in population BR2 evolved in *Ago-2^414/414^*, may represent a case of genetic draft: two synonymous mutations, potentially neutral, linked to a nonsynonymous one that may be the actual target of selection. Yet, the lack of infectious clone for DCV does not allow us to test this hypothesis.

Some of the mutations we found to be associated with positive selection coefficients were synonymous changes (Table 1). However, equating synonymous mutations with neutral mutations in compacted RNA genomes has proved to be misleading^54,55^. Selection operates at different levels of a virus’ life cycle, and not all these levels necessarily depend on the amino acid sequence of encoded proteins. For instance, a lack of matching between virus and host codon usages would slowdown translational speed and efficiency^56^; mutations affecting the folding of regulatory secondary structures at noncoding regions would affect the interaction with host and viral factors and thus impact the expression of downstream genes (*e.g*., mutations 5’UTR/A280U, IGR/A6108G and 3’UTR/U9163A all with significant fitness effects -Table 1)^57^; or evasion from antiviral RNAi defenses by changing the most important relevant sites in the target of siRNAs^12,13^.

It is interesting to observe that viral diversity in mutants for antiviral RNAi, which mode of action relies on a direct interaction with the viral genome, did not display increased diversity when compared to mutants from the other immune pathways. One could expect that the release of the selective pressure that RNAi exerts on the virus genome may allow for the appearance of mutations in the viral suppressor of RNAi. Nonetheless, we did not observed such a change, possibly because the RNAi suppressor in DCV shares the first 99 amino acids of the RdRp^58,59^ and mutations could affect polymerase activity. The antiviral action of the other immune pathways remains still unknown and it might even be indirect, with the known role of Imd, Toll, and Egfr pathways in controlling fly microbiota^37,39^ and possibly affecting the prevalence of virus infections. In this regard, it is important to highlight that the diversity of DCV in the *Dif^1/1^* mutant (Toll pathway, already described not to have an impact on DCV defense^60^), was undistinguishable from *w^1118^*, pointing to the specific - although uncharacterized - antiviral functions of these other immune pathways.

Another consideration when interpreting our results is the nature of the viral stock used. This viral stock has been maintained for years in drosophila S2 cells. The observation that viral population diversity decreased along passages in the fly, highlights the strength of the selection forces that constrain the virus to adapt to a new environment. During the successive passages, in the absence of a given immune response, the capacity of the virus to evolve will be determined by a combination of two factors: the adaptation to the new environment (constrain) and the lack of immune response (relaxation). Because DCV replication is significantly increased in all immune deficient mutants, the potential for population diversification is higher. This effect is clearly observed in *w^1118^* flies where the virus is “only” adapting to the new environment and DCV populations evolved in *w^1118^* flies show less variation than all other lineages. Future experimental evolution studies using viral stocks derived from flies, instead of cell cultures, are warranted to address this topic.

In a parallel study published in this issue, Navarro *et al*. used *Arabidopsis thaliana* and turnip mosaic virus (TuMV) to carry out experimental virus evolution assays with a similar design to ours. In their work, the authors used plant mutants compromised in their antiviral response (more permissive to viral infection) or with an enhanced antiviral response (less permissive to viral infection) and allow the virus to evolve for 12 passages. Similarly to what we found in the *Drosophila melanogaster* - DCV system, the authors showed that viral population evolutions dynamics, as well as viral loads, depend on host genotype. Interestingly, a reduction of ancestral genetic variation regardless of the immune pathway affected was also clearly observed, in agreement with our observations.

Taken together, these two studies point to the concerted action of the different immune pathways to limit viral evolution. Response to infection does not simply consist of activating immune pathways, it also encompasses a broad range of physiological consequences including metabolic adaptations, stress responses and tissue repair. Critically, upon infection, the homeostatic regulation of these pathways is altered. However, such alterations do not always result in increased disease severity or acute infections and can also lead to improved survival (or health) despite active virus replication, which defines tolerance. The Drosophila-DCV arm race seems to be a perfect example of tolerance, and evolution during tolerance remains a field that needs to be studied and described in further detail.

## Materials and Methods

### Fly strains and husbandry

Flies were maintained on a standard cornmeal diet (Bloomington) at a constant temperature of 25 °C. All fly lines were cleaned of possible chronic infections (viruses and Wolbachia) as described previously^61^. The presence or absence of these chronic infections was determined by RT-PCR with specific primers for Nora virus, *Drosophila A virus*, DCV (NoraVfor ATGGCGCCAGTTAGTGCAGACCT, NoraVrev CCTGTTGTTCCAGTTGGGTTCGA DAVfor AGAGTGGCTGTGAGGCAGAT, DAVrev GCCATCTGACAACAGCTTGA, DCVfor GTTGCCTTATCTGCTCTG, DCVrev CGCATAACCATGCTCTTCTG) and by PCR with specific primers *Wolbachia sp* (wspfor TGGTCCAATAAGTGATGAAGAAAC, wsprev AAAAATTAAACGCTACTCCA and wspBfor TTTGCAAGTGAAACAGAAGG, wspBrev GCTTTGCTGGCAAAATGG).

Fly mutant lines for *Dcr-2^L811fsX^* and *Dcr-2^R416X^* ^62^, *Ago-2^414^* ^63^, *Spz^2^* ^64^, *Dif^1^* ^65^, *Rel^E20^* ^66^, *Vago^ΔM10^* ^33^ and *Egfr^t1^* ^67^ were isogenized to *w^1118^* fly line genetic background first by replacing the chromosomes not containing the mutation using balancer chromosomes and then by recombination by backcrossing at least ten times to *w^1118^* line. The presence of the mutation was followed during and at the end of the backcrossing procedure by PCR using specific primers (Dcr2811_3001for TTTGACCCATGACTTTGCGGT, Dcr2811_3294rev CCTTGCAGAGATGCCCCTGTT, Dcr2416_4341for GATTGGCATTACCGTCCCGAA, Dcr2416_4670rev AGCGATTCCTG ATGAGTCTTA, Ago2414_rev TTGTGGATGGCTGTTGTCTCG, Ago251B414_for AGAGTCCCCACTTGAATGGCC, Spz2_for GCCTTTGGCGCTTGCCTAATT, Spz2_rev GCTCCTGCAAAGGAATCGCTC, Dif1_for CTTGGCAATCTTCTCGCACAG, Dif1_rev ATCGTGGTCTCCTGTGTGACG, Rel_Ex4rev AGCTCTCCAGTTTGTGCCGAC, Rel-RD_5′UTRfor CTGGCGTTAGTTTCGGCGTTG, Vagod10_for TTGGCCAACGGAAAGGATGTG, Vagod10_rev TGCCACCGATGATCAATGACA, Egfrt1_for CAAAGCTCGAACCGAAATTA, Egfrt1_rev CTTTCTTAACGTCCACATGA).

### Virus production and titration

S2 DCV stock used to start the experiment was prepared in S2 cells. Cells were maintained in Schneider culture medium and at 25 °C and the appearance of the cells was observed daily. Cells were harvest after when cytopathic effect was detected. The cells were frozen at −80 °C, thawed on ice and centrifuged for 15 min at 15,000 g at 4 °C. The supernatant was recovered, aliquoted and stored at −80 °C. Stocks were titred in S2 cells and titres were measured using the end-point dilution method and expressed as TCID_50_.

To produce the DCV stocks from passages *P* = 1 and *P* = 10 from the evolution experiment half of the population of flies infected with DCV from fly genotype (approx. 250 flies) was homogenized in 1× PBS, homogenates were frozen at −80 °C, then thawed on ice, centrifuged to discard the tissue debris, supernatant was recovered and filtered to discard bacteria contamination, then aliquoted and stored at −80 °C. Stocks were titred in S2 cells and titres were measured using the end-point dilution method and expressed as TCID_50_.

### Viral infections and bacterial infections and survival analysis

To do DCV infections by intrathoracic inoculation, 4 to 5 days old female flies were injected with a Nanoject II apparatus (DrummondScientific) with 50 nl of a viral suspension in 10 mM Tris buffer, pH 7. An injection of the same volume of 10 mM Tris, pH 7 served as a mock-infected control. Infected flies were kept at 25 °C, transferred into fresh vials every 2 days and number of dead flies was scored daily. For the bacterial infections, 4 to 5 days old female flies were intrathoracically injected using a Nanoject II apparatus (DrummondScientific) with 50 nl of the bacterial suspension in 1× PBS buffer, pH 7. An injection of the same volume of 1× PBS buffer served as a mock-infected control. Infected flies were kept at 29 °C, transferred into fresh vials every 2 days and number of dead flies was scored daily.

### Virus experimental evolution

To produce the starting DCV stock (DCV stock) 5 to 6 days old *w^1118^* were intrathoracically injected with 100 TCID_50_ of DCV from a stock produced in S2 drosophila cells (S2 DCV stock) or mock infected. At 4 dpi, *N* = 90 DCV infected flies were placed in cages containing fresh medium, left during 3 days and then removed to place in this DCV contaminated or mock contaminated cages *N* = 500 5 to 6 days old wild type or mutant flies (males and females). Flies were allowed to feed ad libitum during 3 days (oral inoculation period), then moved to a clean cage for 1 day, and further placed into a new clean cage and left during 4 days, when they were harvest (DCV *P* = 1). Contaminated cages were after used to place a new group of flies. This procedure was repeated 10 times (10 DCV Passages, *P* = 1 to *P* = 10) and replicated twice (biological replicates BR1 and BR2).

For statistical analyses, TCID_50_ data were transformed as *T* = log(TCID_50_ + 1) and then fitted to a generalized linear model in which fly genotype *(G)* and BR *(B)* were treated as orthogonal factors. *G* was considered as a fixed effects factor whereas *B* was considered as a random effects factor. Evolutionary passage (*P*) was introduced in the model as a fixed effects covariable. We also considered second and third order interactions between the two factors and the covariable. The model equation thus reads:

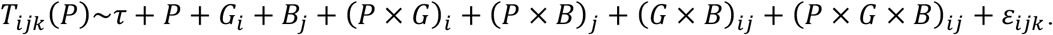

Where *T_ijk_*(*P*) is the transformed TCID_50_ observed for a particular titration assay *k* of BR *j* of fly genotype *i, τ* represents the grand mean value and *ε_ijk_* stands for the error assumed to be Gaussian distributed at every *P*. The significance of each term in the model was evaluated using a likelihood ratio test that follows a *χ*^2^ probability distribution. The magnitude of the effects was evaluated using the 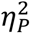 statistic (proportion of total variability in the traits vector attributable to each factor in the model; conventionally, values of 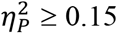 are considered as large effects). These analyses were done using SPSS version 27 (IBM, Armonk, NY).

### RNA extraction, cDNA synthesis and NGS library production

Extraction of RNA from DCV from all passages of the evolution experiment was done using half of the population of infected flies from each fly genotype (approx. 250 flies). The flies were homogenized in Trizol and the manufacturer’s instructions were followed. Total RNA concentration was determined using NanoDrop ND-1000 Spectrophotometer and 300 ng of total RNA were used to produce the cDNA using oligo(dT) as primers to retro-transcription and the Maxima H Minus Reverse Transcriptase Kit, following the manufacturer’s instructions. The cDNA obtained was used as template to amply the full-length genome of DCV with specific primers (DCVfor ATATGTACACACGGCTTTTAGGT and DCVrev CAGTAAGCAGGAAAATTGCG). The PCR products were gel purified and their concentration determined using NanoDrop ND-1000 Spectrophotometer. 200 ng of the purified PCR product were used to produce the NGS library using NEBNext UltraII DNA Library Prep Kit for Illumina and following the manufacturer’s instructions.

Sequencing of DCV populations from *Dif^1/1^* mutant flies from *P* = 4 and *P* = 6 from BR1 and *P* = 8 from BR2 did not work.

### Genetic diversity analyses

#### Variant frequency threshold

To determine the error rate of the sequencing procedure, including library preparation, four sequencing technical replicates from S2 DCV stock were used (Supplementary Figure 3a). First, pairwise comparison was done to identify the variant frequency threshold above which at least 95% of the variants were detected in both considered replicates (highest detection threshold = 0.0028). All variants above detection threshold were then correlated between each technical replicate to ensure good correlation between reported frequency values: the Pearson correlation coefficient between the detected frequency for variants was *r* ≥ 0.982 for all pairwise correlation (*p* < 2.2×10^−16^). The R package used for the analysis has been described elsewhere^68–71^.

#### Nucleotide diversity (π)

Nucleotide diversity of the viral population was computed using the following formula^72^:

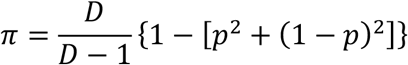

with *D*, the sequencing depth and *p* the frequency of the minority variant at each nucleotide site. For diallelic SNV, π ranges from 0 to 0.5 (both alleles at equal frequency). In the subsequent analyses, π was averaged over all polymorphic nucleotide sites of the DCV genome of each sample^73^. A site was considered polymorphic if at least one sample showed the presence of a nucleotide variant at said position of the DCV genome. Log_10_-transformed site-averaged π values were then compared between fly genotypes (orthogonal factor), biological replicates (orthogonal factor), passages (continuous variable) and genomic regions (orthogonal factor) and their interactions using a generalized linear model. The significance of each term in the model was evaluated using a likelihood ratio test that follows a *χ*^2^ probability distribution.

#### Estimation of relative mutational fitness effects

We have followed the classic population genetics method described in Hartl and Clark (1989)^43^. In short, lets *x_l_(t)* be the frequency of a mutant allele (SNP) at genomic position *l* and passage *t* and, therefore, 1 - *x_l_*(*t*) the frequency of the wild-type allele. It holds that 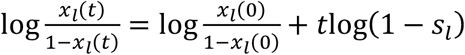, where *s_l_* is the selection coefficient of the mutant relative to the wild-type allele at locus *l*. Selection coefficients calculated this way have units of inverse time (per passage in our case). This equation was fitted to the time series data of each locus *l* shown in Figure 3 by least squares regression, obtaining an estimate of *s_l_* and its standard error (SEM).

Haplotype inference was done using two different statistical approaches. First, by assessing the similarity between temporal dynamics of all possible pairs of loci. To this end, Pearson partial correlation coefficients (controlling for passages) were computed and their significance level corrected for multiple tests of the same null hypothesis using Benjamini and Hochberg (1995)^74^ false discovery rate (FDR) method. Correlation coefficient matrices were visualized as heatmaps in which more similar alleles were clustered together. Second, we confirmed the results from the first method using the longitudinal variant allele frequency factorization problem (LVAFFP) method as implemented in CALDER^75^. LVAFFP generates spanning trees of a directed graph constructed from the variant allele frequencies. The output of CALDER was used as input of TimeScape^76^ to generate the Muller plots that illustrate the ancestry of mutations and haplotypes along the evolution experiment (Figure 3).

Statistical analyses described in this section have been done with R version 4.0.2 in RStudio version 1.3.1073. Scripts are provided in Supplementary File S1.

## Acknowledgements

We thank members of the Saleh Lab and M. Vignuzzi for fruitful discussions. We thank C. Meignin for *Rel^E20^* and *Vago^ΔM10^* flies. This work was supported by the European Research Council (FP7/2013-2019 ERC CoG 615220) and the French Government’s Investissement d’ Avenir program, Laboratoire d’Excellence Integrative Biology of Emerging Infectious Diseases (grant ANR-10-LABX-62-IBEID) to M.-C.S. Work in S.F.E.’s laboratory was supported by grants BFU2015-65037-P and PID2019-103998GB-I00 (Spain Agencia Estatal de Investigación - FEDER) and PROMETEU2019/012 (Generalitat Valenciana).

## Author contributions

V.M. and M.-C.S. conceived the study, and V.M., M.-C.S., A.K. and L.Q.M. established the experimental design. V.M., V.G., and H.B. performed the investigations. S.L. and S.F.E. performed the formal analyses. V.M., S.F.E. and M.-C.S. wrote the paper and acquired funding.

## Competing interests

The authors declare no competing interests.

## Materials & Correspondence

to whom correspondence and material requests should be addressed.

## Notes

### Competing Interest Statement

The authors have declared no competing interest.

