## Supplementary Information for "Innate immune pathways act synergistically to constrain RNA virus evolution in *Drosophila melanogaster*"

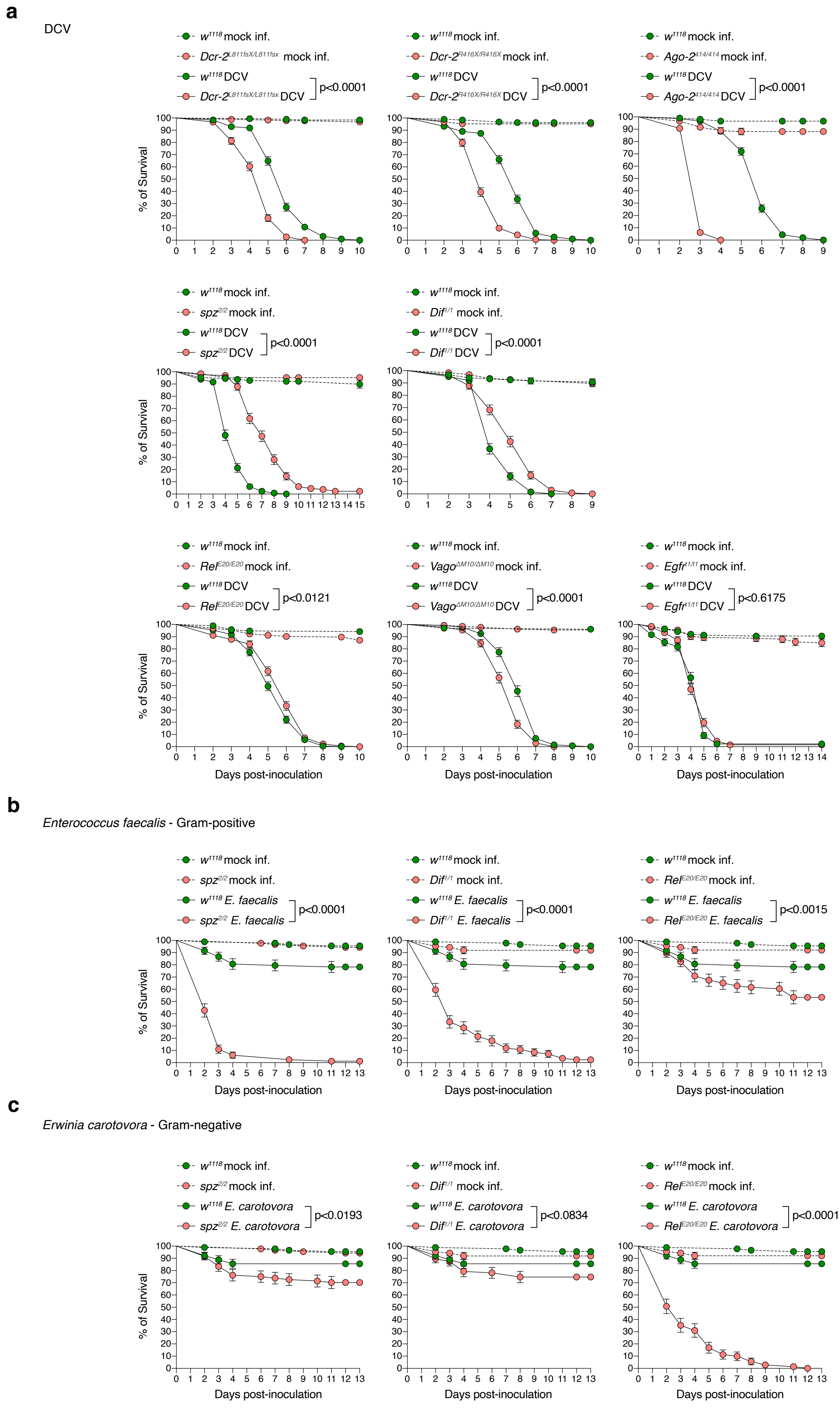


**Supplementary Figure 1. Characterization of the newly produced innate immunity backcrossed fly lines.** Backcrossed innate immunity deficient fly lines were intrathoracically injected with **a)** 10 $TCID_{50}$ units of DCV, **b)** 50 nl of a suspension of optical density (OD) = 10 from *Enterococcus faecalis* (Gram-positive bacteria), and **c)** 50 nl of a suspension of OD = 200 from *E. carotovora carotovora 15* (Ecc15) (Gram-positive bacteria). After viral DCV and *E. faecalis* inoculation flies were kept at 25°C and at 29°C after Ecc15 inoculation. Survival was measured daily. Two independent experiments with three biological replicates of N = 20 flies each per condition were analyzed. Error bars indicate +/-1 SEM; n.s., not significant. Survival curves were compared via log-rank (Mantel–Cox) tests.

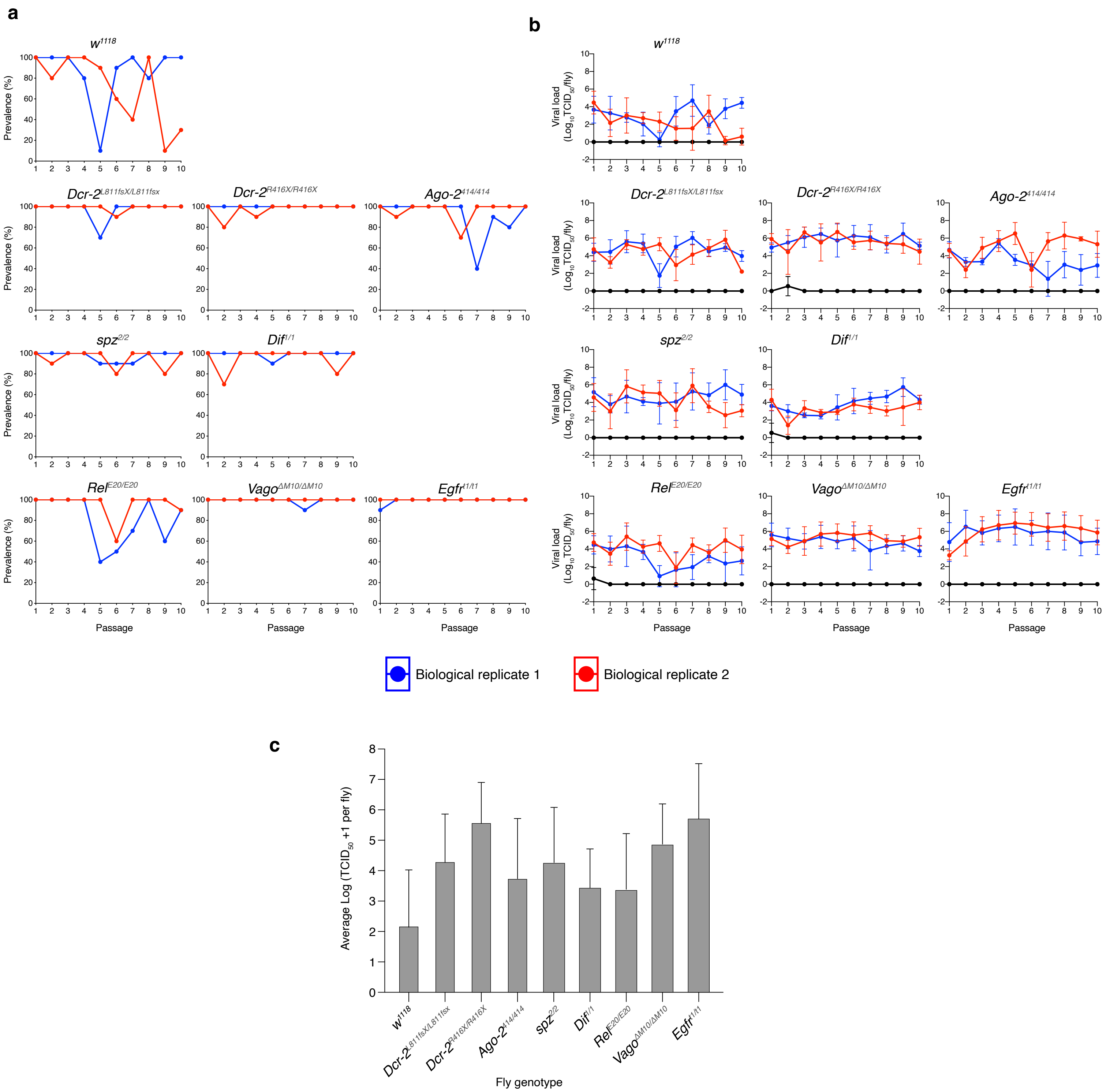


**Supplementary Figure 2. Viral load and prevalence across the DCV evolution experiment.** Viral load of 10 individual flies coming from DCV inoculated cages and four individual flies coming from mock inoculated cages was determined by $TCID_{50}$. **a)** Prevalence, calculated as the percentage of flies positive by $TCID_{50}$. **b)** Viral load in each genotype across the 10 DCV passages. **c)** Average viral loads per individual fly of each genotype estimated from the GLM fitted to the data shown in panel b. Error bars represent +/-1 SD.

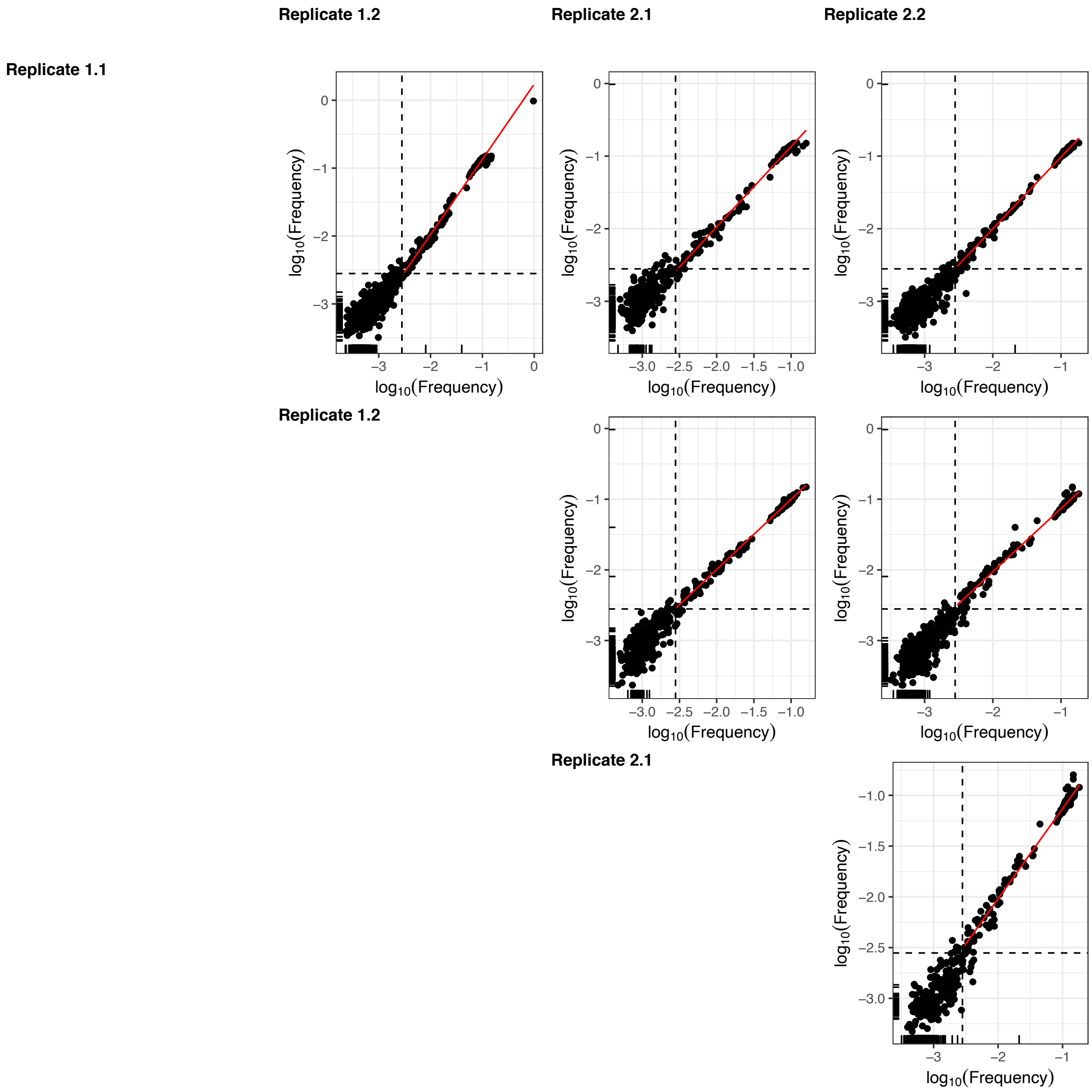


**Supplementary Figure 3. Determination of NGS threshold error and mapping of sequenced derived from DCV starting stocks used.** Pairwise correlation between variant frequency ($\log_{10}$ transformed) in 4 technical sequencing replicates derived from S2 DCV stock. Dashed line represents the frequency threshold value used for subsequent analyses (0.0028). Red line represents the linear regression for variant frequency above the frequency threshold. Black ticks on axis represent missing variants in the other technical replicate under consideration.

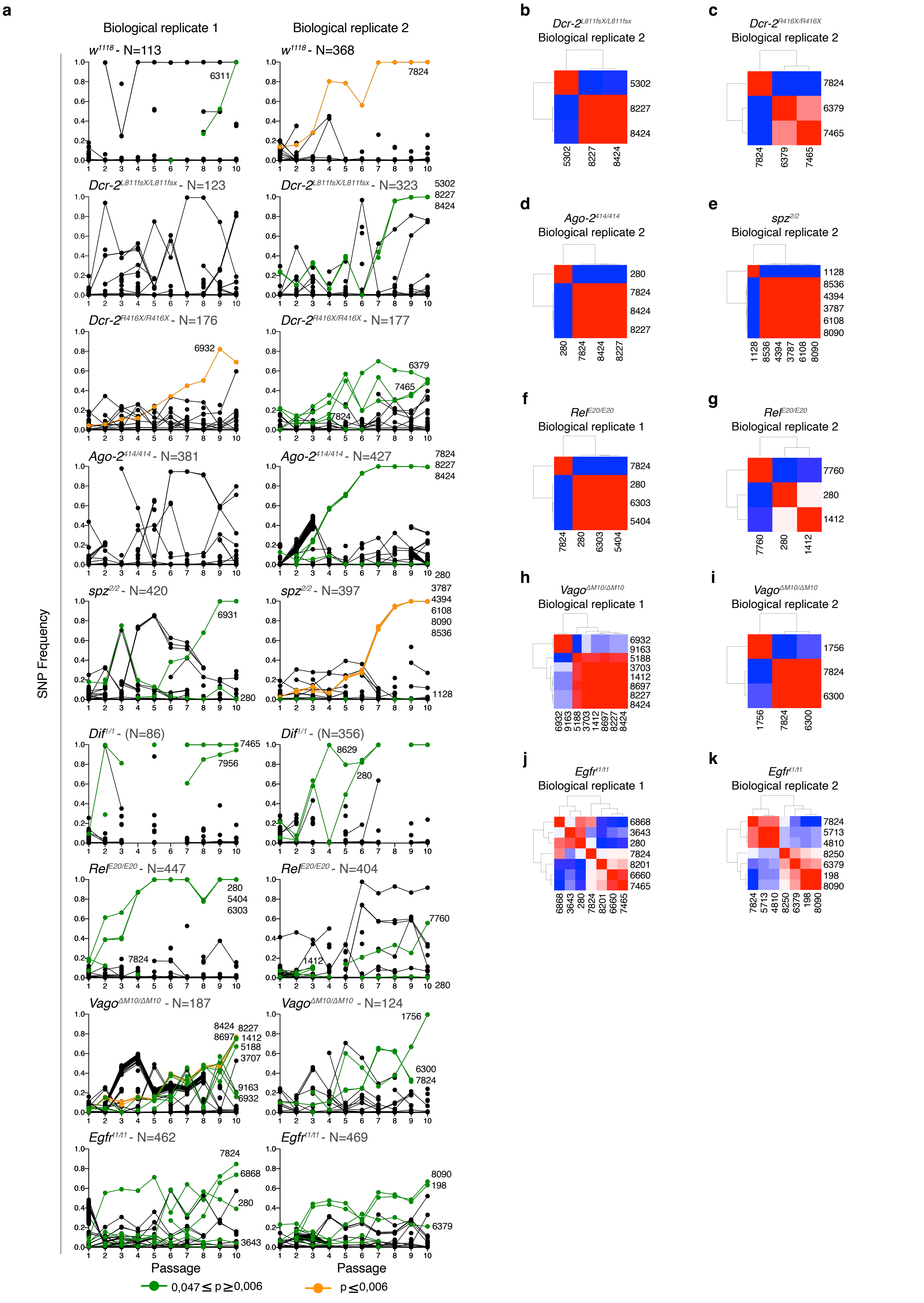


**Supplementary Figure 4. Evolution of DCV variants. a)** Trajectories of DCV variants across passages, **b)** to **k)** pairwise Pearson correlation coefficients between mutations' frequencies along evolutionary time.

**Supplementary Table 1. Pairwise comparisons of viral diversity (π) found in each fly genotype across all passages.**

| | Contrast | Estimate | SE | *d.f.* | *z.ratio* | *p. value* |
|---|---|---|---|---|---|---|
| **Fly genotype** | *Ago-2*$^{414/414}$ - *Dcr-2*$^{R416X/R416X}$ | 0.148531 | 0.124 | Inf | 1.198 | 1.0000 |
| | *Ago-2*$^{414/414}$ - *Dcr-2*$^{L811fsX/L811fsX}$ | 0.330554 | 0.124 | Inf | 2.666 | 0.2762 |
| | *Ago-2*$^{414/414}$ - *Dif*$^{1/1}$ | 0.520045 | 0.130 | Inf | 4.012 | 0.0022 |
| | *Ago-2*$^{414/414}$ - *Egfr*$^{t1/t1}$ | -0.186983 | 0.124 | Inf | -1.508 | 1.0000 |
| | *Ago-2*$^{414/414}$ - *Rel*$^{E20/20}$ | 0.214099 | 0.124 | Inf | 1.727 | 1.0000 |
| | *Ago-2*$^{414/414}$ - *spz*$^{2/2}$ | 0.182274 | 0.124 | Inf | 1.470 | 1.0000 |
| | *Ago-2*$^{414/414}$ - *Vago*$^{DM10/DM10}$ | -0.187586 | 0.124 | Inf | -1.513 | 1.0000 |
| | *Ago-2*$^{414/414}$ - *w*$^{1118}$ | 0.657285 | 0.124 | Inf | 5.301 | <0.0001 |
| | *Dcr-2*$^{R416X/R416X}$ - *Dcr-2*$^{L811fsX/L811fsX}$ | 0.182023 | 0.124 | Inf | 1.468 | 1.0000 |
| | *Dcr-2*$^{R416X/R416X}$ - *Dif*$^{1/1}$ | 0.371515 | 0.130 | Inf | 2.866 | 0.1497 |
| | *Dcr-2*$^{R416X/R416X}$ - *Egfr*$^{t1/t1}$ | -0.335514 | 0.124 | Inf | -2.706 | 0.2451 |
| | *Dcr-2*$^{R416X/R416X}$ - *Rel*$^{E20/20}$ | 0.065568 | 0.124 | Inf | 0.529 | 1.0000 |
| | *Dcr-2*$^{R416X/R416X}$ - *spz*$^{2/2}$ | 0.033743 | 0.124 | Inf | 0.272 | 1.0000 |
| | *Dcr-2*$^{R416X/R416X}$ - *Vago*$^{DM10/DM10}$ | -0.336116 | 0.124 | Inf | -2.711 | 0.2415 |
| | *Dcr-2*$^{R416X/R416X}$ - *w*$^{1118}$ | 0.508755 | 0.124 | Inf | 4.103 | 0.0015 |
| | *Dcr-2*$^{L811fsX/L811fsX}$ - *Dif*$^{1/1}$ | 0.189491 | 0.130 | Inf | 1.462 | 1.0000 |
| | *Dcr-2*$^{L811fsX/L811fsX}$ - *Egfr*$^{t1/t1}$ | -0.517537 | 0.124 | Inf | -4.174 | 0.0011 |
| | *Dcr-2*$^{L811fsX/L811fsX}$ - *Rel*$^{E20/20}$ | -0.116455 | 0.124 | Inf | -0.939 | 1.0000 |
| | *Dcr-2*$^{L811fsX/L811fsX}$ - *spz*$^{2/2}$ | -0.148280 | 0.124 | Inf | -1.196 | 1.0000 |
| | *Dcr-2*$^{L811fsX/L811fsX}$ - *Vago*$^{DM10/DM10}$ | -0.518139 | 0.124 | Inf | -4.179 | 0.0011 |
| | *Dcr-2*$^{L811fsX/L811fsX}$ - *w*$^{1118}$ | 0.326731 | 0.124 | Inf | 2.635 | 0.3026 |
| | *Dif*$^{1/1}$ - *Egfr*$^{t1/t1}$ | -0.707028 | 0.130 | Inf | -5.454 | <0.0001 |
| | *Dif*$^{1/1}$ - *Rel*$^{E20/20}$ | -0.305946 | 0.130 | Inf | -2.360 | 0.6577 |
| | *Dif*$^{1/1}$ - *spz*$^{2/2}$ | -0.337772 | 0.130 | Inf | -2.606 | 0.3302 |
| | *Dif*$^{1/1}$ - *Vago*$^{DM10/DM10}$ | -0.707631 | 0.130 | Inf | -5.459 | <0.0001 |
| | *Dif*$^{1/1}$ - *w*$^{1118}$ | 0.137240 | 0.130 | Inf | 1.059 | 1.0000 |
| | *Egfr*$^{t1/t1}$ - *Rel*$^{E20/20}$ | 0.401082 | 0.124 | Inf | 3.235 | 0.0438 |
| | *Egfr*$^{t1/t1}$ - *spz*$^{2/2}$ | 0.369257 | 0.124 | Inf | 2.978 | 0.1043 |
| | *Egfr*$^{t1/t1}$ - *Vago*$^{DM10/DM10}$ | -0.000602 | 0.124 | Inf | -0.005 | 1.0000 |
| | *Egfr*$^{t1/t1}$ - *w*$^{1118}$ | 0.844268 | 0.124 | Inf | 6.810 | <0.0001 |
| | *Rel*$^{E20/20}$ - *spz*$^{2/2}$ | -0.031825 | 0.124 | Inf | -0.257 | 1.0000 |
| | *Rel*$^{E20/20}$ - *Vago*$^{DM10/DM10}$ | -0.401685 | 0.124 | Inf | -3.240 | 0.0431 |
| | *Rel*$^{E20/20}$ - *w*$^{1118}$ | 0.443186 | 0.124 | Inf | 3.575 | 0.0126 |
| | *spz*$^{2/2}$ - *Vago*$^{DM10/DM10}$ | -0.369859 | 0.124 | Inf | -2.983 | 0.1027 |
| | *spz*$^{2/2}$ - *w*$^{1118}$ | 0.475012 | 0.124 | Inf | 3.831 | 0.0046 |

| | | | | | | |
|---|---|---|---|---|---|---|
| | *Vago*$^{DM10/DM10}$ - *w*$^{1118}$ | 0.844871 | 0.124 | Inf | 6.814 | <0.0001 |
| **Genome region** | 3'UTR - 5'UTR IRES | -0.876 | 0.0645 | Inf | -13.577 | <0.0001 |
| | 3'UTR - ORF1 | -1.348 | 0.0645 | Inf | -20.915 | <0.0001 |
| | 3'UTR - ORF2 | -1.473 | 0.0645 | Inf | -22.860 | <0.0001 |
| | 5'UTR IRES - ORF1 | -0.472 | 0.0512 | Inf | -9.214 | <0.0001 |
| | 5'UTR IRES - ORF2 | -0.597 | 0.0512 | Inf | -11.660 | <0.0001 |
| | ORF1 - ORF2 | -0.125 | 0.0512 | Inf | -2.450 | 0.0857 |
| **Fly genotype – P1** | *Ago-2*$^{414/414}$ - *Dcr-2*$^{R416X/R416X}$ | 0.55657 | 0.604 | 10 | 0.922 | 1.0000 |
| | *Ago-2*$^{414/414}$ - *Dcr-2*$^{L811fsX/L811fsX}$ | 0.69861 | 0.604 | 10 | 1.157 | 1.0000 |
| | *Ago-2*$^{414/414}$ - S2 DCV stock R1 | -0.70848 | 0.604 | 10 | -1.173 | 1.0000 |
| | *Ago-2*$^{414/414}$ - S2 DCV stock R2 | -0.71263 | 0.604 | 10 | -1.180 | 1.0000 |
| | *Ago-2*$^{414/414}$ - *Dif*$^{1/1}$ | -0.02819 | 0.604 | 10 | -0.047 | 1.0000 |
| | *Ago-2*$^{414/414}$ - *Egfr*$^{t1/t1}$ | -0.26877 | 0.604 | 10 | -0.445 | 1.0000 |
| | *Ago-2*$^{414/414}$ - *Rel*$^{E20/20}$ | -0.29062 | 0.604 | 10 | -0.481 | 1.0000 |
| | *Ago-2*$^{414/414}$ - *spz*$^{2/2}$ | -0.06647 | 0.604 | 10 | -0.110 | 1.0000 |
| | *Ago-2*$^{414/414}$ - *Vago*$^{DM10/DM10}$ | 0.55166 | 0.604 | 10 | 0.914 | 1.0000 |
| | *Ago-2*$^{414/414}$ - *w*$^{1118}$ | 0.01750 | 0.604 | 10 | 0.029 | 1.0000 |
| | *Dcr-2*$^{R416X/R416X}$ - *Dcr-2*$^{L811fsX/L811fsX}$ | 0.14204 | 0.604 | 10 | 0.235 | 1.0000 |
| | *Dcr-2*$^{R416X/R416X}$ - S2 DCV stock R1 | -1.26505 | 0.604 | 10 | -2.095 | 1.0000 |
| | *Dcr-2*$^{R416X/R416X}$ - S2 DCV stock R2 | -1.26920 | 0.604 | 10 | -2.102 | 1.0000 |
| | *Dcr-2*$^{R416X/R416X}$ - *Dif*$^{1/1}$ | -0.58476 | 0.604 | 10 | -0.968 | 1.0000 |
| | *Dcr-2*$^{R416X/R416X}$ - *Egfr*$^{t1/t1}$ | -0.82534 | 0.604 | 10 | -1.367 | 1.0000 |
| | *Dcr-2*$^{R416X/R416X}$ - *Rel*$^{E20/20}$ | -0.84719 | 0.604 | 10 | -1.403 | 1.0000 |
| | *Dcr-2*$^{R416X/R416X}$ - *spz*$^{2/2}$ | -0.62304 | 0.604 | 10 | -1.032 | 1.0000 |
| | *Dcr-2*$^{R416X/R416X}$ - *Vago*$^{DM10/DM10}$ | -0.00491 | 0.604 | 10 | -0.008 | 1.0000 |
| | *Dcr-2*$^{R416X/R416X}$ - *w*$^{1118}$ | -0.53907 | 0.604 | 10 | -0.893 | 1.0000 |
| | *Dcr-2*$^{L811fsX/L811fsX}$ - S2 DCV stock R1 | -1.40709 | 0.604 | 10 | -2.330 | 1.0000 |
| | *Dcr-2*$^{L811fsX/L811fsX}$ - S2 DCV stock R2 | -1.41124 | 0.604 | 10 | -2.337 | 1.0000 |
| | *Dcr-2*$^{L811fsX/L811fsX}$ - *Dif*$^{1/1}$ | -0.72680 | 0.604 | 10 | -1.204 | 1.0000 |
| | *Dcr-2*$^{L811fsX/L811fsX}$ - *Egfr*$^{t1/t1}$ | -0.96738 | 0.604 | 10 | -1.602 | 1.0000 |
| | *Dcr-2*$^{L811fsX/L811fsX}$ - *Rel*$^{E20/20}$ | -0.98923 | 0.604 | 10 | -1.638 | 1.0000 |
| | *Dcr-2*$^{L811fsX/L811fsX}$ - *spz*$^{2/2}$ | -0.76508 | 0.604 | 10 | -1.267 | 1.0000 |
| | *Dcr-2*$^{L811fsX/L811fsX}$ - *Vago*$^{DM10/DM10}$ | -0.14695 | 0.604 | 10 | -0.243 | 1.0000 |
| | *Dcr-2*$^{L811fsX/L811fsX}$ - *w*$^{1118}$ | -0.68111 | 0.604 | 10 | -1.128 | 1.0000 |
| | S2 DCV stock R1 - S2 DCV stock R2 | -0.00415 | 0.604 | 10 | -0.007 | 1.0000 |
| | S2 DCV stock R1- *Dif*$^{1/1}$ | 0.68029 | 0.604 | 10 | 1.127 | 1.0000 |
| | S2 DCV stock R1- *Egfr*$^{t1/t1}$ | 0.43971 | 0.604 | 10 | 0.728 | 1.0000 |
| | S2 DCV stock R1- *Rel*$^{E20/20}$ | 0.41786 | 0.604 | 10 | 0.692 | 1.0000 |

| | | | | | | |
|---|---|---|---|---|---|---|
| | S2 DCV stock R1 - *spz*$^{2/2}$ | 0.64201 | 0.604 | 10 | 1.063 | 1.0000 |
| | S2 DCV stock R1- *Vago*$^{DM10/DM10}$ | 1.26014 | 0.604 | 10 | 2.087 | 1.0000 |
| | S2 DCV stock R1- *w*$^{1118}$ | 0.72598 | 0.604 | 10 | 1.202 | 1.0000 |
| | S2 DCV stock R2- *Dif*$^{1/1}$ | 0.68444 | 0.604 | 10 | 1.133 | 1.0000 |
| | S2 DCV stock R2- *Egfr*$^{t1/t1}$ | 0.44386 | 0.604 | 10 | 0.735 | 1.0000 |
| | S2 DCV stock R2- *Rel*$^{E20/20}$ | 0.42201 | 0.604 | 10 | 0.699 | 1.0000 |
| | S2 DCV stock R2 - *spz*$^{2/2}$ | 0.64616 | 0.604 | 10 | 1.070 | 1.0000 |
| | S2 DCV stock R2- *Vago*$^{DM10/DM10}$ | 1.26429 | 0.604 | 10 | 2.094 | 1.0000 |
| | S2 DCV stock R2- *w*$^{1118}$ | 0.73013 | 0.604 | 10 | 1.209 | 1.0000 |
| | *Dif*$^{1/1}$ - *Egfr*$^{t1/t1}$ | -0.24058 | 0.604 | 10 | -0.398 | 1.0000 |
| | *Dif*$^{1/1}$ - *Rel*$^{E20/20}$ | -0.26243 | 0.604 | 10 | -0.435 | 1.0000 |
| | *Dif*$^{1/1}$ - *spz*$^{2/2}$ | -0.03828 | 0.604 | 10 | -0.063 | 1.0000 |
| | *Dif*$^{1/1}$ - *Vago*$^{DM10/DM10}$ | 0.57985 | 0.604 | 10 | 0.960 | 1.0000 |
| | *Dif*$^{1/1}$ - *w*$^{1118}$ | 0.04569 | 0.604 | 10 | 0.076 | 1.0000 |
| | *Egfr*$^{t1/t1}$ - *Rel*$^{E20/20}$ | -0.02185 | 0.604 | 10 | -0.036 | 1.0000 |
| | *Egfr*$^{t1/t1}$ - *spz*$^{2/2}$ | 0.20230 | 0.604 | 10 | 0.335 | 1.0000 |
| | *Egfr*$^{t1/t1}$ - *Vago*$^{DM10/DM10}$ | 0.82043 | 0.604 | 10 | 1.359 | 1.0000 |
| | *Egfr*$^{t1/t1}$ - *w*$^{1118}$ | 0.28627 | 0.604 | 10 | 0.474 | 1.0000 |
| | *Rel*$^{E20/20}$ - *spz*$^{2/2}$ | 0.22415 | 0.604 | 10 | 0.371 | 1.0000 |
| | *Rel*$^{E20/20}$ - *Vago*$^{DM10/DM10}$ | 0.84228 | 0.604 | 10 | 1.395 | 1.0000 |
| | *Rel*$^{E20/20}$ - *w*$^{1118}$ | 0.30812 | 0.604 | 10 | 0.510 | 1.0000 |
| | *spz*$^{2/2}$ - *Vago*$^{DM10/DM10}$ | 0.61813 | 0.604 | 10 | 1.024 | 1.0000 |
| | *spz*$^{2/2}$ - *w*$^{1118}$ | 0.08397 | 0.604 | 10 | 0.139 | 1.0000 |
| | *Vago*$^{DM10/DM10}$ - *w*$^{1118}$ | -0.53416 | 0.604 | 10 | -0.885 | 1.0000 |
| **Fly genotype - P5** | *Ago-2*$^{414/414}$ - *Dcr-2*$^{R416X/R416X}$ | -0.02796 | 0.29 | 10 | -0.096 | 1.0000 |
| | *Ago-2*$^{414/414}$ - *Dcr-2*$^{L811fsX/L811fsX}$ | -0.08041 | 0.29 | 10 | -0.277 | 1.0000 |
| | *Ago-2*$^{414/414}$ - S2 DCV stock R1 | -1.29970 | 0.29 | 10 | -4.481 | 0.0647 |
| | *Ago-2*$^{414/414}$ - S2 DCV stock R2 | -1.30385 | 0.29 | 10 | -4.495 | 0.0633 |
| | *Ago-2*$^{414/414}$ - *Dif*$^{1/1}$ | 0.10898 | 0.29 | 10 | 0.376 | 1.0000 |
| | *Ago-2*$^{414/414}$ - *Egfr*$^{t1/t1}$ | -0.15550 | 0.29 | 10 | -0.536 | 1.0000 |
| | *Ago-2*$^{414/414}$ - *Rel*$^{E20/20}$ | 0.31647 | 0.29 | 10 | 1.091 | 1.0000 |
| | *Ago-2*$^{414/414}$ - *spz*$^{2/2}$ | 0.01315 | 0.29 | 10 | 0.045 | 1.0000 |
| | *Ago-2*$^{414/414}$ - *Vago*$^{DM10/DM10}$ | -0.44370 | 0.29 | 10 | -1.530 | 1.0000 |
| | *Ago-2*$^{414/414}$ - *w*$^{1118}$ | 0.56139 | 0.29 | 10 | 1.936 | 1.0000 |
| | *Dcr-2*$^{R416X/R416X}$ - *Dcr-2*$^{L811fsX/L811fsX}$ | -0.05245 | 0.29 | 10 | -0.181 | 1.0000 |
| | *Dcr-2*$^{R416X/R416X}$ - S2 DCV stock R1 | -1.27174 | 0.29 | 10 | -4.385 | 0.0752 |
| | *Dcr-2*$^{R416X/R416X}$ - S2 DCV stock R2 | -1.27589 | 0.29 | 10 | -4.399 | 0.0736 |
| | *Dcr-2*$^{R416X/R416X}$ - *Dif*$^{1/1}$ | 0.13694 | 0.29 | 10 | 0.472 | 1.0000 |

| | | | | | |
|---|---|---|---|---|---|
| *Dcr-2*$^{R416X/R416X}$ - *Egfr*$^{t1/t1}$ | -0.12754 | 0.29 | 10 | -0.440 | 1.0000 |
| *Dcr-2*$^{R416X/R416X}$ - *Rel*$^{E20/20}$ | 0.34443 | 0.29 | 10 | 1.187 | 1.0000 |
| *Dcr-2*$^{R416X/R416X}$ - *spz*$^{2/2}$ | 0.04111 | 0.29 | 10 | 0.142 | 1.0000 |
| *Dcr-2*$^{R416X/R416X}$ - *Vago*$^{DM10/DM10}$ | -0.41574 | 0.29 | 10 | -1.433 | 1.0000 |
| *Dcr-2*$^{R416X/R416X}$ - *w*$^{1118}$ | 0.58935 | 0.29 | 10 | 2.032 | 1.0000 |
| *Dcr-2*$^{L811fsX/L811fsX}$ - S2 DCV stock R1 | -1.21929 | 0.29 | 10 | -4.204 | 1.0000 |
| *Dcr-2*$^{L811fsX/L811fsX}$ - S2 DCV stock R2 | -1.22345 | 0.29 | 10 | -4.218 | 0.0977 |
| *Dcr-2*$^{L811fsX/L811fsX}$ - *Dif*$^{1/1}$ | 0.18939 | 0.29 | 10 | 0.653 | 1.0000 |
| *Dcr-2*$^{L811fsX/L811fsX}$ - *Egfr*$^{t1/t1}$ | -0.07510 | 0.29 | 10 | -0.259 | 1.0000 |
| *Dcr-2*$^{L811fsX/L811fsX}$ - *Rel*$^{E20/20}$ | 0.39688 | 0.29 | 10 | 1.368 | 1.0000 |
| *Dcr-2*$^{L811fsX/L811fsX}$ - *spz*$^{2/2}$ | 0.09356 | 0.29 | 10 | 0.323 | 1.0000 |
| *Dcr-2*$^{L811fsX/L811fsX}$ - *Vago*$^{DM10/DM10}$ | -0.36330 | 0.29 | 10 | -1.253 | 1.0000 |
| *Dcr-2*$^{L811fsX/L811fsX}$ - *w*$^{1118}$ | 0.64180 | 0.29 | 10 | 2.213 | 1.0000 |
| S2 DCV stock R1 - S2 DCV stock R2 | -0.00415 | 0.29 | 10 | -0.014 | 1.0000 |
| S2 DCV stock R1- *Dif*$^{1/1}$ | 1.40868 | 0.29 | 10 | 4.857 | 0.0366 |
| S2 DCV stock R1- *Egfr*$^{t1/t1}$ | 1.14420 | 0.29 | 10 | 3.945 | 0.1514 |
| S2 DCV stock R1- *Rel*$^{E20/20}$ | 1.61617 | 0.29 | 10 | 5.572 | 0.0130 |
| S2 DCV stock R1 - *spz*$^{2/2}$ | 1.31285 | 0.29 | 10 | 4.526 | 0.0604 |
| S2 DCV Stock R1- *Vago*$^{DM10/DM10}$ | 0.85600 | 0.29 | 10 | 2.951 | 0.7977 |
| S2 DCV stock R1- *w*$^{1118}$ | 1.86109 | 0.29 | 10 | 6.416 | 0.0042 |
| S2 DCV stock R2- *Dif*$^{1/1}$ | 1.41283 | 0.29 | 10 | 4.871 | 0.0358 |
| S2 DCV stock R2- *Egfr*$^{t1/t1}$ | 1.14835 | 0.29 | 10 | 3.959 | 0.1480 |
| S2 DCV stock R2- *Rel*$^{E20/20}$ | 1.62032 | 0.29 | 10 | 5.586 | 0.0128 |
| S2 DCV stock R2 - *spz*$^{2/2}$ | 1.31701 | 0.29 | 10 | 4.541 | 0.0591 |
| S2 DCV stock R2- *Vago*$^{DM10/DM10}$ | 0.86015 | 0.29 | 10 | 2.966 | 0.7784 |
| S2 DCV stock R2- *w*$^{1118}$ | 1.86524 | 0.29 | 10 | 6.431 | 0.0041 |
| *Dif*$^{1/1}$ - *Egfr*$^{t1/t1}$ | -0.26449 | 0.29 | 10 | -0.912 | 1.0000 |
| *Dif*$^{1/1}$ - *Rel*$^{E20/20}$ | 0.20749 | 0.29 | 10 | 0.715 | 1.0000 |
| *Dif*$^{1/1}$ - *spz*$^{2/2}$ | -0.09583 | 0.29 | 10 | 0.330 | 1.0000 |
| *Dif*$^{1/1}$ - *Vago*$^{DM10/DM10}$ | -0.55268 | 0.29 | 10 | -1.905 | 1.0000 |
| *Dif*$^{1/1}$ - *w*$^{1118}$ | 0.45241 | 0.29 | 10 | 1.560 | 1.0000 |
| *Egfr*$^{t1/t1}$ - *Rel*$^{E20/20}$ | 0.47198 | 0.29 | 10 | 1.627 | 1.0000 |
| *Egfr*$^{t1/t1}$ - *spz*$^{2/2}$ | 0.16866 | 0.29 | 10 | 0.581 | 1.0000 |
| *Egfr*$^{t1/t1}$ - *Vago*$^{DM10/DM10}$ | -0.28820 | 0.29 | 10 | -0.994 | 1.0000 |
| *Egfr*$^{t1/t1}$ - *w*$^{1118}$ | 0.71690 | 0.29 | 10 | 2.472 | 1.0000 |
| *Rel*$^{E20/20}$ - *spz*$^{2/2}$ | -0.30332 | 0.29 | 10 | -1.046 | 1.0000 |
| *Rel*$^{E20/20}$ - *Vago*$^{DM10/DM10}$ | -0.76017 | 0.29 | 10 | -2.621 | 1.0000 |
| *Rel*$^{E20/20}$ - *w*$^{1118}$ | 0.24492 | 0.29 | 10 | 0.844 | 1.0000 |

| | | | | | | |
|---|---|---|---|---|---|---|
| | $spz^{2/2}$ - $Vago^{DM10/DM10}$ | -0.45686 | 0.29 | 10 | -1.575 | 1.0000 |
| | $spz^{2/2}$ - $w^{1118}$ | 0.54824 | 0.29 | 10 | 1.890 | 1.0000 |
| | $Vago^{DM10/DM10}$ - $w^{1118}$ | 1.00509 | 0.29 | 10 | 3.465 | 0.3338 |
| **Fly genotype - P10** | $Ago\text{-}2^{414/414}$ - $Dcr\text{-}2^{R416X/R416X}$ | 0.00110 | 0.245 | 10 | 0.004 | 1.0000 |
| | $Ago\text{-}2^{414/414}$ - $Dcr\text{-}2^{L811fsX/L811fsX}$ | 0.33463 | 0.245 | 10 | 1.364 | 1.0000 |
| | $Ago\text{-}2^{414/414}$ - S2 DCV stock R1 | -1.10416 | 0.245 | 10 | -4.501 | 0.0628 |
| | $Ago\text{-}2^{414/414}$ - S2 DCV stock R2 | -1.10831 | 0.245 | 10 | -4.517 | 0.0612 |
| | $Ago\text{-}2^{414/414}$ - $Dif^{1/1}$ | 0.83303 | 0.245 | 10 | 3.395 | 0.3753 |
| | $Ago\text{-}2^{414/414}$ - $Egfr^{t1/t1}$ | 0.06986 | 0.245 | 10 | 0.285 | 1.0000 |
| | $Ago\text{-}2^{414/414}$ - $Rel^{E20/20}$ | 0.43982 | 0.245 | 10 | 1.793 | 1.0000 |
| | $Ago\text{-}2^{414/414}$ - $spz^{2/2}$ | 0.57632 | 0.245 | 10 | 2.349 | 1.0000 |
| | $Ago\text{-}2^{414/414}$ - $Vago^{DM10/DM10}$ | 0.09496 | 0.245 | 10 | 0.387 | 1.0000 |
| | $Ago\text{-}2^{414/414}$ - $w^{1118}$ | 0.49881 | 0.245 | 10 | 2.033 | 1.0000 |
| | $Dcr\text{-}2^{R416X/R416X}$ - $Dcr\text{-}2^{L811fsX/L811fsX}$ | 0.33353 | 0.245 | 10 | 1.359 | 1.0000 |
| | $Dcr\text{-}2^{R416X/R416X}$ - S2 DCV stock R1 | -1.10526 | 0.245 | 10 | -4.505 | 0.0624 |
| | $Dcr\text{-}2^{R416X/R416X}$ - S2 DCV stock R2 | -1.10941 | 0.245 | 10 | -4.522 | 0.0608 |
| | $Dcr\text{-}2^{R416X/R416X}$ - $Dif^{1/1}$ | 0.83193 | 0.245 | 10 | 3.391 | 0.3781 |
| | $Dcr\text{-}2^{R416X/R416X}$ - $Egfr^{t1/t1}$ | 0.06876 | 0.245 | 10 | 0.280 | 1.0000 |
| | $Dcr\text{-}2^{R416X/R416X}$ - $Rel^{E20/20}$ | 0.43872 | 0.245 | 10 | 1.788 | 1.0000 |
| | $Dcr\text{-}2^{R416X/R416X}$ - $spz^{2/2}$ | 0.57522 | 0.245 | 10 | 2.345 | 1.0000 |
| | $Dcr\text{-}2^{R416X/R416X}$ - $Vago^{DM10/DM10}$ | 0.09386 | 0.245 | 10 | 0.383 | 1.0000 |
| | $Dcr\text{-}2^{R416X/R416X}$ - $w^{1118}$ | 0.49771 | 0.245 | 10 | 2.029 | 1.0000 |
| | $Dcr\text{-}2^{L811fsX/L811fsX}$ - S2 DCV stock R1 | -1.43879 | 0.245 | 10 | -5.864 | 0.0087 |
| | $Dcr\text{-}2^{L811fsX/L811fsX}$ - S2 DCV stock R2 | -1.44294 | 0.245 | 10 | -5.881 | 0.0085 |
| | $Dcr\text{-}2^{L811fsX/L811fsX}$ - $Dif^{1/1}$ | 0.49841 | 0.245 | 10 | 2.032 | 1.0000 |
| | $Dcr\text{-}2^{L811fsX/L811fsX}$ - $Egfr^{t1/t1}$ | -0.26477 | 0.245 | 10 | -1.079 | 1.0000 |
| | $Dcr\text{-}2^{L811fsX/L811fsX}$ - $Rel^{E20/20}$ | 0.10519 | 0.245 | 10 | 0.429 | 1.0000 |
| | $Dcr\text{-}2^{L811fsX/L811fsX}$ - $spz^{2/2}$ | 0.24169 | 0.245 | 10 | 0.985 | 1.0000 |
| | $Dcr\text{-}2^{L811fsX/L811fsX}$ - $Vago^{DM10/DM10}$ | -0.23966 | 0.245 | 10 | -0.977 | 1.0000 |
| | $Dcr\text{-}2^{L811fsX/L811fsX}$ - $w^{1118}$ | 0.16419 | 0.245 | 10 | 0.669 | 1.0000 |
| | S2 DCV stock R1 - S2 DCV stock R2 | -0.00415 | 0.245 | 10 | -0.017 | 1.0000 |
| | S2 DCV stock R1- $Dif^{1/1}$ | 1.93719 | 0.245 | 10 | 7.896 | 0.0007 |
| | S2 DCV stock R1- $Egfr^{t1/t1}$ | 1.17402 | 0.245 | 10 | 4.785 | 0.0407 |
| | S2 DCV stock R1- $Rel^{E20/20}$ | 1.54398 | 0.245 | 10 | 6.293 | 0.0049 |
| | S2 DCV stock R1 - $spz^{2/2}$ | 1.68048 | 0.245 | 10 | 6.850 | 0.0025 |
| | S2 DCV stock R1- $Vago^{DM10/DM10}$ | 1.19912 | 0.245 | 10 | 4.888 | 0.0349 |
| | S2 DCV stock R1- $w^{1118}$ | 1.60297 | 0.245 | 10 | 6.534 | 0.0036 |
| | S2 DCV stock R2- $Dif^{1/1}$ | 1.94134 | 0.245 | 10 | 7.913 | 0.0007 |

| | | | | | |
|---|---|---|---|---|---|
| S2 DCV stock R2- *Egfr*$^{t1/t1}$ | 1.17817 | 0.245 | 10 | 4.802 | 0.0397 |
| S2 DCV stock R2- *Rel*$^{E20/20}$ | 1.54813 | 0.245 | 10 | 6.310 | 0.0048 |
| S2 DCV stock R2 - *spz*$^{2/2}$ | 1.68463 | 0.245 | 10 | 6.867 | 0.0024 |
| S2 DCV stock R2- *Vago*$^{DM10/DM10}$ | 1.20327 | 0.245 | 10 | 4.905 | 0.0340 |
| S2 DCV stock R2- *w*$^{1118}$ | 1.60713 | 0.245 | 10 | 6.551 | 0.0036 |
| *Dif*$^{1/1}$ - *Egfr*$^{t1/t1}$ | -0.76318 | 0.245 | 10 | -3.111 | 0.6076 |
| *Dif*$^{1/1}$ - *Rel*$^{E20/20}$ | -0.39322 | 0.245 | 10 | -1.603 | 1.0000 |
| *Dif*$^{1/1}$ - *spz*$^{2/2}$ | -0.25671 | 0.245 | 10 | 1.046 | 1.0000 |
| *Dif*$^{1/1}$ - *Vago*$^{DM10/DM10}$ | -0.73807 | 0.245 | 10 | -3.008 | 0.7235 |
| *Dif*$^{1/1}$ - *w*$^{1118}$ | -0.33422 | 0.245 | 10 | -1.362 | 1.0000 |
| *Egfr*$^{t1/t1}$ - *Rel*$^{E20/20}$ | 0.36996 | 0.245 | 10 | 1.508 | 1.0000 |
| *Egfr*$^{t1/t1}$ - *spz*$^{2/2}$ | 0.50646 | 0.245 | 10 | 2.064 | 1.0000 |
| *Egfr*$^{t1/t1}$ - *Vago*$^{DM10/DM10}$ | 0.02510 | 0.245 | 10 | 0.102 | 1.0000 |
| *Egfr*$^{t1/t1}$ - *w*$^{1118}$ | 0.42896 | 0.245 | 10 | 1.748 | 1.0000 |
| *Rel*$^{E20/20}$ - *spz*$^{2/2}$ | 0.13650 | 0.245 | 10 | 0.556 | 1.0000 |
| *Rel*$^{E20/20}$ - *Vago*$^{DM10/DM10}$ | -0.34486 | 0.245 | 10 | -1.406 | 1.0000 |
| *Rel*$^{E20/20}$ - *w*$^{1118}$ | 0.05900 | 0.245 | 10 | 0.240 | 1.0000 |
| *spz*$^{2/2}$ - *Vago*$^{DM10/DM10}$ | -0.48136 | 0.245 | 10 | 1.962 | 1.0000 |
| *spz*$^{2/2}$ - *w*$^{1118}$ | -0.07751 | 0.245 | 10 | -0.316 | 1.0000 |
| *Vago*$^{DM10/DM10}$ - *w*$^{1118}$ | 0.40385 | 0.245 | 10 | 1.646 | 1.0000 |

Two biological replicates and two technical replicates were performed from S2 DCV stock. For the purpose of this analysis each biological replicate was pooled, S2 DCV stock R1 and R2 respectively.

### Supplementary Table 2. Statistical analysis of the survival curves from Figure 5 - Log-rank (Mantel–Cox) tests

**Viral Passage 1 – Biological replicate 1**

| Viral stock Origin | Nr. of flies | Median survival | Viral stock Origin | | | | | | | | | | |
|---|---|---|---|---|---|---|---|---|---|---|---|---|---|
| | | | **Mock** | **S2 DCV stock** | **DCV stock** | ***w*^1118** | ***Dcr-2*^L811fsX/L811fsX** | ***Dcr-2*^R416X/R416X** | ***Ago-2*^414/414** | ***spz*^2/2** | ***Dif*^1/1** | ***Rel*^E20/20** | ***Vago*^DM10/DM10** |
| **Mock** | 235 | Und. | | | | | | | | | | | |
| **S2 DCV stock** | 235 | 5 | <0,0001 | | | | | | | | | | |
| **DCV stock** | 231 | 6 | <0,0001<br>**** | <0,0001<br>**** | | | | | | | | | |
| ***w*^1118** | 119 | 5 | <0,0001<br>**** | 0,1152<br>ns | <0,0001<br>**** | | | | | | | | |
| ***Dcr-2*^L811fsX/L811fsX** | 120 | 4 | <0,0001<br>**** | <0,0001<br>**** | <0,0001<br>**** | <0,0001<br>**** | | | | | | | |
| ***Dcr-2*^R416X/R416X** | 118 | 5 | <0,0001<br>**** | 0,3054<br>ns | <0,0001<br>**** | 0,6783<br>ns | <0,0001<br>**** | | | | | | |
| ***Ago-2*^414/414** | 120 | 4 | <0,0001<br>**** | <0,0001<br>**** | <0,0001<br>**** | <0,0001<br>**** | 0,1412<br>ns | <0,0001<br>**** | | | | | |
| ***spz*^2/2** | 119 | 5 | <0,0001<br>**** | 0,2037<br>ns | <0,0001<br>**** | 0,0007<br>*** | <0,0001<br>**** | 0,0061<br>** | <0,0001<br>**** | | | | |
| ***Dif*^1/1** | 117 | 5 | <0,0001<br>**** | 0,0888<br>ns | <0,0001<br>**** | 0,8372<br>ns | <0,0001<br>**** | 0,7587<br>ns | <0,0001<br>**** | 0,0004<br>*** | | | |
| ***Rel*^E20/20** | 118 | 5 | <0,0001<br>**** | <0,0001<br>**** | <0,0001<br>**** | 0,0222<br>* | <0,0001<br>**** | 0,0024<br>** | 0,0007<br>*** | <0,0001<br>**** | 0,0044<br>** | | |
| ***Vago*^DM10/DM10** | 120 | 5 | <0,0001<br>**** | 0,0038<br>** | <0,0001<br>**** | 0,3019<br>ns | <0,0001<br>**** | 0,1366<br>ns | 0,0004<br>*** | <0,0001<br>**** | 0,1696<br>ns | 0,3443<br>ns | |
| ***Egfr*^t1/t1** | 119 | 6 | <0,0001<br>**** | <0,0001<br>**** | 0,0002<br>*** | <0,0001<br>**** | <0,0001<br>**** | <0,0001<br>**** | <0,0001<br>**** | 0,0038<br>** | <0,0001<br>**** | <0,0001<br>**** | <0,0001<br>**** |

**Viral Passage 10 – Biological replicate 1**

| Viral stock Origin | Nr. of flies | Median survival | Viral stock Origin | | | | | | | | | | |
|---|---|---|---|---|---|---|---|---|---|---|---|---|---|
| | | | **Mock** | **S2 DCV stock** | **DCV stock** | ***w*^1118** | ***Dcr-2*^L811fsX/L811fsX** | ***Dcr-2*^R416X/R416X** | ***Ago-2*^414/414** | ***spz*^2/2** | ***Dif*^1/1** | ***Rel*^E20/20** | ***Vago*^DM10/DM10** |
| **Mock** | 235 | Und. | | | | | | | | | | | |
| **S2 DCV stock** | 235 | 5 | <0,0001 | | | | | | | | | | |
| **DCV stock** | 231 | 6 | <0,0001<br>**** | <0,0001<br>**** | | | | | | | | | |

| | | | | | | | | | | | | | |
|---|---|---|---|---|---|---|---|---|---|---|---|---|---|
| ***w*$^{1118}$** | 116 | 6 | <0,0001 **** | 0,3626 ns | <0,0001 **** | | | | | | | | |
| ***Dcr-2*$^{L811fsX/L811fsX}$** | 114 | 6 | <0,0001 **** | <0,0001 **** | 0,0655 *ns* | <0,0001 **** | | | | | | | |
| ***Dcr-2*$^{R416X/R416X}$** | 106 | 6 | <0,0001 **** | 0,0008 *** | <0,0001 **** | 0,0228 * | 0,0029 ** | | | | | | |
| ***Ago-2*$^{414/414}$** | 115 | 6 | <0,0001 **** | <0,0001 **** | 0,5588 ns | <0,0001 **** | 0,1497 ns | 0,0003 *** | | | | | |
| ***spz*$^{2/2}$** | 118 | 5 | <0,0001 **** | 0,353 ns | <0,0001 **** | 0,6788 ns | <0,0001 **** | 0,004 ** | <0,0001 **** | | | | |
| ***Dif*$^{1/1}$** | 117 | 6 | <0,0001 **** | 0,0063 ** | <0,0001 **** | 0,1699 ns | <0,0001 **** | 0,2717 ns | <0,0001 **** | 0,0416 * | | | |
| ***Rel*$^{E20/20}$** | 110 | 6 | <0,0001 **** | <0,0001 **** | 0,0020 ** | 0,0027 ** | 0,2122 ns | 0,2453 ns | 0,0185 * | 0,0014 ** | 0,0493 * | | |
| ***Vago*$^{DM10/DM10}$** | 108 | 5 | <0,0001 **** | 0,0126 * | <0,0001 **** | 0,147 ns | 0,0003 *** | 0,428 ns | <0,0001 **** | 0,074 ns | 0,9034 ns | 0,066 ns | |
| ***Egfr*$^{t1/t1}$** | 115 | 6 | <0,0001 **** | <0,0001 **** | 0,8472 ns | <0,0001 **** | 0,017 * | <0,0001 **** | 0,607 ns | <0,0001 **** | <0,0001 **** | 0,0011 ** | <0,0001 **** |

**Viral Passage 1 – Biological replicate 2**

| **Viral stock Origin** | Nr. of flies | **Median survival** | **Viral stock Origin** | | | | | | | | | | |
|---|---|---|---|---|---|---|---|---|---|---|---|---|---|
| | | | **Mock** | **S2 DCV stock** | **DCV stock** | ***w*$^{1118}$** | ***Dcr-2*$^{L811fsX/L811fsX}$** | ***Dcr-2*$^{R416X/R416X}$** | ***Ago-2*$^{414/414}$** | ***spz*$^{2/2}$** | ***Dif*$^{1/1}$** | ***Rel*$^{E20/20}$** | ***Vago*$^{DM10/DM10}$** |
| **Mock** | 235 | Und. | | | | | | | | | | | |
| **S2 DCV stock** | 225 | 5 | <0,0001 **** | | | | | | | | | | |
| **DCV stock** | 233 | 6 | <0,0001 **** | <0,0001 **** | | | | | | | | | |
| ***w*$^{1118}$** | 119 | 4 | <0,0001 **** | <0,0001 **** | <0,0001 **** | | | | | | | | |
| ***Dcr-2*$^{L811fsX/L811fsX}$** | 119 | 5 | <0,0001 **** | 0,0209 * | <0,0001 **** | 0,0300 * | | | | | | | |
| ***Dcr-2*$^{R416X/R416X}$** | 119 | 5 | <0,0001 **** | 0,2219 ns | <0,0001 **** | <0,0001 **** | 0,0012 ** | | | | | | |
| ***Ago-2*$^{414/414}$** | 118 | 4 | <0,0001 **** | <0,0001 **** | <0,0001 **** | 0,3835 ns | 0,0025 ** | <0,0001 **** | | | | | |
| ***spz*$^{2/2}$** | 119 | 5 | <0,0001 **** | 0,2456 ns | <0,0001 **** | <0,0001 **** | 0,0011 ** | 0,9665 ns | <0,0001 **** | | | | |
| ***Dif*$^{1/1}$** | 119 | 4 | <0,0001 **** | <0,0001 **** | <0,0001 **** | 0,2122 ns | 0,3402 ns | <0,0001 **** | 0,0329 * | <0,0001 **** | | | |

| | | | | | | | | | | | | | |
|---|---|---|---|---|---|---|---|---|---|---|---|---|---|
| ***Rel***$^{E20/20}$ | 120 | 5 | <0,0001 **** | <0,0001 **** | <0,0001 **** | 0,009 ** | 0,6047 ns | 0,0062 ** | 0,0005 *** | 0,005 ** | 0,1455 ns | | |
| ***Vago***$^{DM10/DM10}$ | 120 | 6 | <0,0001 **** | <0,0001 **** | <0,0001 **** | <0,0001 **** | <0,0001 **** | <0,0001 **** | <0,0001 **** | <0,0001 **** | <0,0001 **** | <0,0001 **** | |
| ***Egfr***$^{t1/t1}$ | 115 | 5 | <0,0001 **** | 0,1738 ns | 0,5815 ns | <0,0001 **** | 0,0005 *** | 0,1995 ns | <0,0001 **** | 0,1971 ns | <0,0001 **** | 0,0016 ** | 0,0033 ** |

**Viral Passage 10 – Biological replicate 2**

| **Viral stock Origin** | **Nr. of flies** | **Median survival** | **Viral stock Origin** | | | | | | | | | | |
|---|---|---|---|---|---|---|---|---|---|---|---|---|---|
| | | | **Mock** | **S2 DCV stock** | **DCV stock** | ***w***$^{1118}$ | ***Dcr-2***$^{L811fsX/L811fsX}$ | ***Dcr-2***$^{R416X/R416X}$ | ***Ago-2***$^{414/414}$ | ***spz***$^{2/2}$ | ***Dif***$^{1/1}$ | ***Rel***$^{E20/20}$ | ***Vago***$^{DM10/DM10}$ |
| **Mock** | 235 | Und. | | | | | | | | | | | |
| **S2 DCV stock** | 225 | 5 | <0,0001 **** | | | | | | | | | | |
| **DCV stock** | 233 | 6 | <0,0001 **** | <0,0001 **** | | | | | | | | | |
| ***w***$^{1118}$ | 104 | 5 | <0,0001 **** | 0,0209 * | 0,0083 ** | | | | | | | | |
| ***Dcr-2***$^{L811fsX/L811fsX}$ | 105 | 5 | <0,0001 **** | 0,1381 ns | 0,0004 *** | 0,3873 ns | | | | | | | |
| ***Dcr-2***$^{R416X/R416X}$ | 110 | 5 | <0,0001 **** | 0,0011 ** | 0,0211 * | 0,6294 ns | 0,1213 ns | | | | | | |
| ***Ago-2***$^{414/414}$ | 111 | 5 | <0,0001 **** | 0,1376 ns | 0,0005 *** | 0,4395 ns | 0,9246 ns | 0,1763 ns | | | | | |
| ***spz***$^{2/2}$ | 102 | 5 | <0,0001 **** | 0,0003 *** | 0,2429 ns | 0,2615 ns | 0,0400 * | 0,3852 ns | 0,0602 ns | | | | |
| ***Dif***$^{1/1}$ | 98 | 5 | <0,0001 **** | 0,0979 ns | 0,0018 ** | 0,5727 ns | 0,698 ns | 0,3219 ns | 0,8399 ns | 0,1019 ns | | | |
| ***Rel***$^{E20/20}$ | 100 | 5 | <0,0001 **** | 0,0905 ns | 0,002 ** | 0,6142 ns | 0,7401 ns | 0,3066 ns | 0,7993 ns | 0,1219 ns | 0,9375 ns | | |
| ***Vago***$^{DM10/DM10}$ | 94 | 5 | <0,0001 **** | 0,0013 ** | 0,1121 ns | 0,3741 ns | 0,0779 ns | 0,5576 ns | 0,0992 ns | 0,8418 ns | 0,146 ns | 0,1781 ns | |
| ***Egfr***$^{t1/t1}$ | 98 | 5 | <0,0001 **** | 0,0297 * | 0,0101 * | 0,9411 ns | 0,4563 ns | 0,5765 ns | 0,5522 ns | 0,1878 ns | 0,7472 ns | 0,7471 ns | 0,3464 ns |
